# Wetting-mediated extracellular phase separation drives long-range cell adhesion

**DOI:** 10.64898/2026.01.23.700778

**Authors:** Anheng Wang, Dixi Yang, Haijiao Zhang, Vesselin Paunov, Shuo Tian, Lei Dong, Hajime Tanaka, Jiaxing Yuan, Chunming Wang

**Affiliations:** Institute of Chinese Medical Sciences & State Key Laboratory of Mechanism and Quality of Chinese Medicine, University of Macau, Macau SAR 999078, China; Department of Pharmaceutical Sciences, Faculty of Health Sciences, University of Macau, Macau SAR 999078, China; Zhuhai UM Science and Technology Research Institute, University of Macau, Hengqin 519031, Guangdong, China; Advanced Materials Thrust, Function Hub, The Hong Kong University of Science and Technology (Guangzhou), Nansha District, Guangzhou 511453, China; National Center for Protein Science, Chinese Academy of Sciences, Shanghai 200031, China; Department of Chemistry, School of Sciences and Humanities, Nazarbayev University, Kabanbay Baryr 53, Astana 010000, Kazakhstan; Guangzhou Doublle Bioproduct Co., Ltd.,Guangzhou,510535, China; Chemistry and Biomedicine Innovation Center, Nanjing University, Nanjing, Jiangsu 210023, China; Research Center for Advanced Science and Technology, University of Tokyo, 4-6-1 Komaba, Meguro-ku, Tokyo 153-8904, Japan; Department of Fundamental Engineering, Institute of Industrial Science, University of Tokyo, 4-6-1 Komaba, Meguro-ku, Tokyo 153-8505, Japan

## Abstract

Cells must efficiently locate and engage for tissue formation and immune coordination, yet classical receptor-ligand binding is limited to nanometre distances and is inherently slow [1–3]. Here, we uncover a previously unrecognised physical principle, liquid-like adhesion by phase separation (LAPS). This process creates dynamic wetting layers on cell surfaces [4, 5], functioning as “liquid bridges” that enable robust, long-range cell capture across tens of micrometres. Remarkably, this wetting-mediated attraction remains effective at nanomolar concentrations—conditions where bulk phase separation would not be expected—and facilitates high-fidelity cell sorting through competitive wetting. By integrating aqueous two-phase systems, endogenous proteins (Galectin-3, CCL5), and fluid-particle-dynamics simulations [6], we demonstrate that extracellular liquid-liquid phase separation not only mediates long-range cell capture but also acts as a physical catalyst for contact-dependent signaling. These findings establish extracellular phase separation as a key physical principle complementing molecular recognition in multicellular systems, offering new opportunities for understanding immune response, tissue morphogenesis, and therapeutic strategies targeting the extracellular environment [7, 8].

## INTRODUCTION

The efficiency of cellular life depends on orchestrating biochemical reactions that are powerfully regulated liquid-liquid phase separation (LLPS), which creates transient, membraneless organelles [9–11]. However, the role of LLPS extends beyond mere chemical sequestration. It is increasingly understood that intracellular condensates are also active physical players, generating capillary-like forces and interfacial tensions that mechanically sculpt and organise the cell’s internal architecture [12–17]. This establishes LLPS as a fundamental engine for physical self-organisation on a subcellular scale. This raises a crucial and largely unexplored question: what happens when this physical engine is unleashed from the confines of a single cell into the vast extracellular space? The principle dictates that if LLPS can generate organizing forces between organelles, its effects should be magnified when operating between cells, escalating to the macroscopic scale of tissue organisation. While recent studies have indeed begun to report the existence of functional extracellular LLPS [18–24], the new physical laws that emerge when this mechanism operates in an intercellular regime remain largely unknown.

Answering this question is crucial, as our understanding of intercellular interaction has been dominated by a “chemical paradigm”: the binding of individual receptor-ligand pairs, a model of discrete, short-range “molecular bonds” [1]. While this picture is essential for explaining biochemical specificity, it is supplemented only by generic physical forces, such as van der Waals and electrostatic interactions, which are fundamentally weak and short-ranged under physiological conditions [25–27]. This “molecular bond” worldview is therefore insufficient to explain rapid, long-range coordinations over several microns required to assemble tissues or orchestrate immune responses.

We propose that multicellular organisation arises not from the discrete chemistry of individual bonds, but from a distinct physical mechanism: the “collective thermodynamics” of the extracellular milieu, mediated specifically by phase-separated condensates on cell surfaces. We term this new mechanism of interaction Liquid-like Adhesion by Phase Separation (LAPS). We hypothesise that LAPS is achieved via “surface wetting”, where the cell membrane acts as a catalytic surface to nucleate a condensed liquid phase even under bulk one-phase concentration [28, 29]. The emergence of this new liquid phase, we propose, fundamentally transforms the cell-environment interface from a conventional two-phase “cell–liquid” model into a dynamic three-phase “cell–liquid–liquid” system [4, 5, 30]. The critical consequence of this transformation is the emergence of attractive intercellular forces that drive long-range cell adhesion, as if the cells generate their own hands.

In this work, we combine cell-based experiments, endogenous biomolecules, and Fluid Particle Dynamics (FPD) simulations to systematically deconstruct this new physical paradigm. We directly measure these emergent forces, demonstrate how they can be harnessed to perform high-fidelity cell sorting, and show how the resulting liquid bridges can act as physical catalysts for signaling. Our findings establish a new, general physical principle underlying multicellularity, opening up a new field that investigates the continuum physics of cell-cell interactions. This new paradigm is neutral to specific biological outcomes, providing a physical foundation upon which the diverse functional consequences of extracellular phase separation—both beneficial and pathological—can be explored by future studies.

## RESULTS AND DISCUSSION

### Morphological Transition of Cell Populations

To investigate how LAPS reshapes cellular organisation, we first characterised the transformation of a simple two-phase “cell–liquid” suspension into a three-phase “cell–liquid– liquid” environment. To systematically map this morphological phase space, we examined the behaviour of Jurkat cells in a canonical Dextran (DEX)/Polyethylene Glycol (PEG) aqueous two-phase system (ATPS).

Our experiments revealed that cells display a rich spectrum of morphologies —ranging from pendular to capillary states — depending on the relative composition of the surrounding phases (Fig. 1a–c). The cell volume fraction was fixed at *ϕ*_Cell_ = 0.01 throughout. Without DEX (*ϕ*_DEX_ = 0), cells remained fully dispersed. Introducing a small DEX fraction (*ϕ*_DEX_ = 0.01) rapidly induced aggregation, producing a dynamic, pendular network characteristic of capillary bridging (Fig. 1b). Concave DEX menisci formed between adjacent cells, confirming preferential wetting of the cell surface by DEX over PEG. Notably, the number of clusters peaked at 5 minutes, and more than 80% of cells aggregated within ~3 minutes (Fig. S1a, Video 1). In sharp contrast, aggregation in a conventional single-phase system — mediated by integrins [31] —required nearly 2 hours (Fig. S1b) and won’t aggregate within 30 mins in 5wt.% PEG solution (Video 2).

**FIG. 1.**
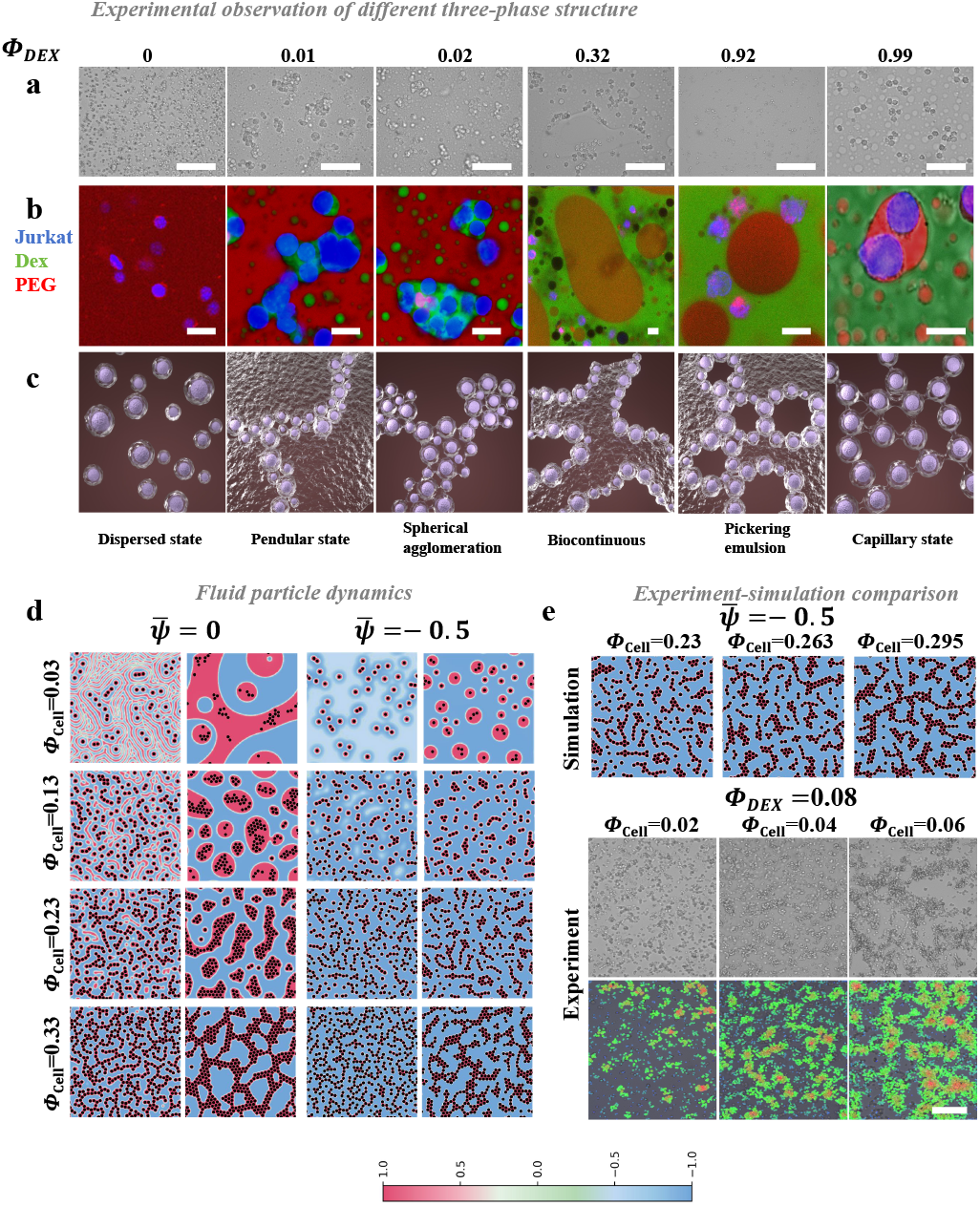
Morphological transitions of cellular aggregates governed by wetting effects and phase volume. **a** Bright-field images showing the systematic morphological transition of Jurkat cells in a PEG/DEX aqueous two-phase system (ATPS) as a function of the DEX volume fraction (*ϕ*_DEX_). Scale bar, 50 *µ*m. **b** Confocal microscopy confirming preferential wetting of Jurkat cells (blue, Cytotell Blue) by the DEX-rich phase (green, FITC–DEX), suspended within the PEG-rich phase (red, Cy5–PEG). Scale bar, 10 *µ*m. **c** Representative experimentally observed states: Dispersed, Pendular, Spherical agglomeration, Bicontinuous, Pickering emulsion, and Capillary states. **d** State diagram from two-dimensional FPD simulations, mapping morphological states as a function of cell occupation fraction (*ϕ*_Cell_) and system composition 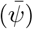 at *t* = 50*τ*_0_ and *t* = 5000*τ*_0_. **e** Direct comparison of simulation and experiment. Top: simulations at fixed 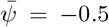 with increasing particle volume fractions. Bottom: experiments at fixed *ϕ*_DEX_ = 0.08 with increasing cell volume fractions (*ϕ*_Cell_ = 0.02, 0.04, 0.06). Scale bar, 50 *µ*m.

As the DEX volume fraction, *ϕ*_DEX_, increased (*ϕ*_DEX_ = 0.02 and above), the system transitioned to spherical agglomerates, as the DEX phase became sufficient to fully wet cells. At near-symmetric composition (*ϕ*_DEX_ ≈ 0.32), cells localised at the liquid–liquid interface, producing a bicontinuous structure. In DEX-dominant systems (*ϕ*_DEX_ = 0.92), cells adsorbed to PEG droplet surfaces, acting as Pickering stabilisers [32, 33] and generating a Pickering emulsion with PEG as the dispersed phase. Notably, at very high DEX content (*ϕ*_DEX_ = 0.99), the system reverted to a classic capillary state: PEG formed convex menisci on cell surfaces, consistent with its weaker wettability relative to DEX [34–36].

Together, these results demonstrate that cellular morphology within ATPS is tightly governed by wetting interactions and phase composition, establishing a physical framework for how LAPS drives rapid, diverse, and reversible modes of cell self-organisation.

The results of two-dimensional (2D) simulations are summarised in Fig. 1d, where we systematically varied the average composition 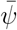 — related to the minority phase fraction via 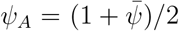 — and the cell occupation fraction *ϕ*_cell_. Consistent with experiments, particles with preferential affinity for one component rapidly developed wetting layers on their surfaces, leading to eventual incorporation into the more wettable phase. The final aggregate morphology was critically dependent on the interplay between *ϕ*_cell_ and 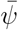 (Fig. 1d).

At symmetric composition (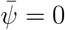, Fig. 1d, left), the system exhibited a striking morphological sequence with increasing *ϕ*_cell_. At low particle fractions (*ϕ*_cell_ = 0.03), cells formed an interconnected network resembling a bicontinuous structure. As *ϕ*_cell_ increased (0.13–0.22), this network fragmented into rounded droplets, mimicking spherical agglomeration. At the highest concentration (*ϕ*_cell_ = 0.33), the system re-entered a capillary-like state characterised by network-forming aggregates.

In contrast, at asymmetric composition (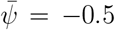, Fig. 1d, right), the morphological pathway differed substantially. At low *ϕ*_cell_, isolated spherical aggregates formed readily (Video 3). As concentration increased, these aggregates rapidly coalesced into denser clusters and eventually transitioned into a capillary-like network (Video 4).

We further explored intermediate regimes by varying 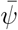 between −0.4 and −0.1 (Figs. S2– S9). Combined with Fig. 1d, these data highlight a key morphological trend: increasing compositional asymmetry promotes branched, fractal-like clusters rather than compact spherical aggregates. Moreover, the re-entrant network–droplet–network transition with increasing *ϕ*_cell_ was unique to the symmetric case 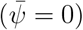.

Together, these simulations reproduce the morphological diversity observed in our experiments and establish that wetting-driven self-organisation is governed by a subtle coupling between composition and cell density.

Quantitative analysis of coarsening dynamics using the average wavenumber ⟨*q*⟩ (Figs. S2– S9) confirmed that domain growth is suppressed at late stages. In all conditions, the presence of particles markedly accelerated early-stage coarsening due to wetting effects, but arrested the subsequent kinetics compared to particle-free systems (*ϕ*_Cell_ = 0) [37]. The dependence of the coarsening exponent on both 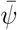 and *ϕ*_Cell_ highlights a subtle coupling between thermodynamic driving forces and the mechanical frustration introduced by particle aggregates [4, 38].

We next compared simulations with experimental results for the asymmetric case 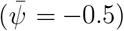 (Fig. 1e). The simulations reflected the experimental trend, with cells assembling into progressively larger and denser clusters as *ϕ*_Cell_ increased. The close correspondence between predicted morphologies and observed patterns strongly supports that our model captures the essential physics of wetting-driven cellular self-organisation.

By integrating experimental and computational approaches, we show that cells in a phase-separating fluid can self-organise into a wide spectrum of morphologies —including branched pendular states, spherical agglomerates, network structures, and emulsions — determined jointly by system composition and cell occupation fraction. A particularly striking observation is the paradoxical enhancement of aggregation as the volume of the high-affinity wetting phase decreases. In this regime, the scarcity of the minority phase drives its condensation into tighter, more robust capillary bridges, thereby enforcing intimate cell–cell contact. This behaviour closely parallels the formation of biological tight junctions and suggests a generic physical mechanism that may facilitate rapid intercellular signal transduction [39, 40].

### Cellular Aggregation in Bulk Binodal and One-Phase regimes

Previous experiments and simulations revealed dramatic morphological transitions, with inter-particle connections forming even when the minority phase was scarce (e.g., *ϕ*_DEX_ ≈ 0.01). A key consideration, however, is the absolute molecular concentration. For example, a 1 v/v% phase of 5 wt.% 500 kDa dextran corresponds to a molar concentration of approximately 1 *µ*M, which is near the threshold for bulk phase separation (**Extended Data Fig**. 1a). This value is substantially higher than the concentrations of most biological signaling factors, which typically act in the nanomolar (nM) range. Thus, a central question is whether capillary bridging remains effective at such physiologically relevant low concentrations, where bulk phase separation is not expected.

To address this, stock solutions of 500 kDa DEX were prepared at various weight percentages and mixed with 5 wt.% PEG at different *ϕ*_DEX_ to yield working concentrations spanning 50–1000 nM, thereby covering both the bulk one-phase and binodal regimes. Remarkably, robust, well-defined cellular networks still formed at bulk one phase regime (500nM DEX) (**Extended Data Fig**. 1b). As the concentration decreased, network integrity gradually weakened, though significant aggregation persisted even at 100 nM (Video 5). A fully dispersed state was observed only below ~50 nM (**Extended Data Fig**. 1b), indicating that even extremely dilute conditions can still support measurable aggregation.

In the nominal one-phase regime (100–500 nM), we observed a wetting behaviour distinct from complete surface coverage. Instead of forming continuous layers, DEX assembled into discrete, punctate aggregates on the cell surfaces, a morphology characteristic of partial wetting due to the scarcity of the wetting phase (Fig. 2a). Despite this minimal coverage, the localised wetting was still sufficient to mediate robust cell–cell attraction. We quantified the aggregation kinetics by tracking the average number of neighbors (*N*_neigh_) per cell over time (Fig. 2b). As expected, the kinetics were concentration-dependent; for instance, aggregation at 100 nM required nearly 10 minutes to plateau, compared to 5 minutes at 1000 nM, though both are dramatically faster than conventional integrin-mediated aggregation [31]. It is important to note that the eventual decline in *N*_neigh_ after its peak does not indicate disaggregation, but is an algorithmic artifact that occurs as interior cells in dense, fully assembled networks become difficult to track. Thus, the peak in *N*_neigh_ signifies the point of maximum network assembly.

**FIG. 2.**
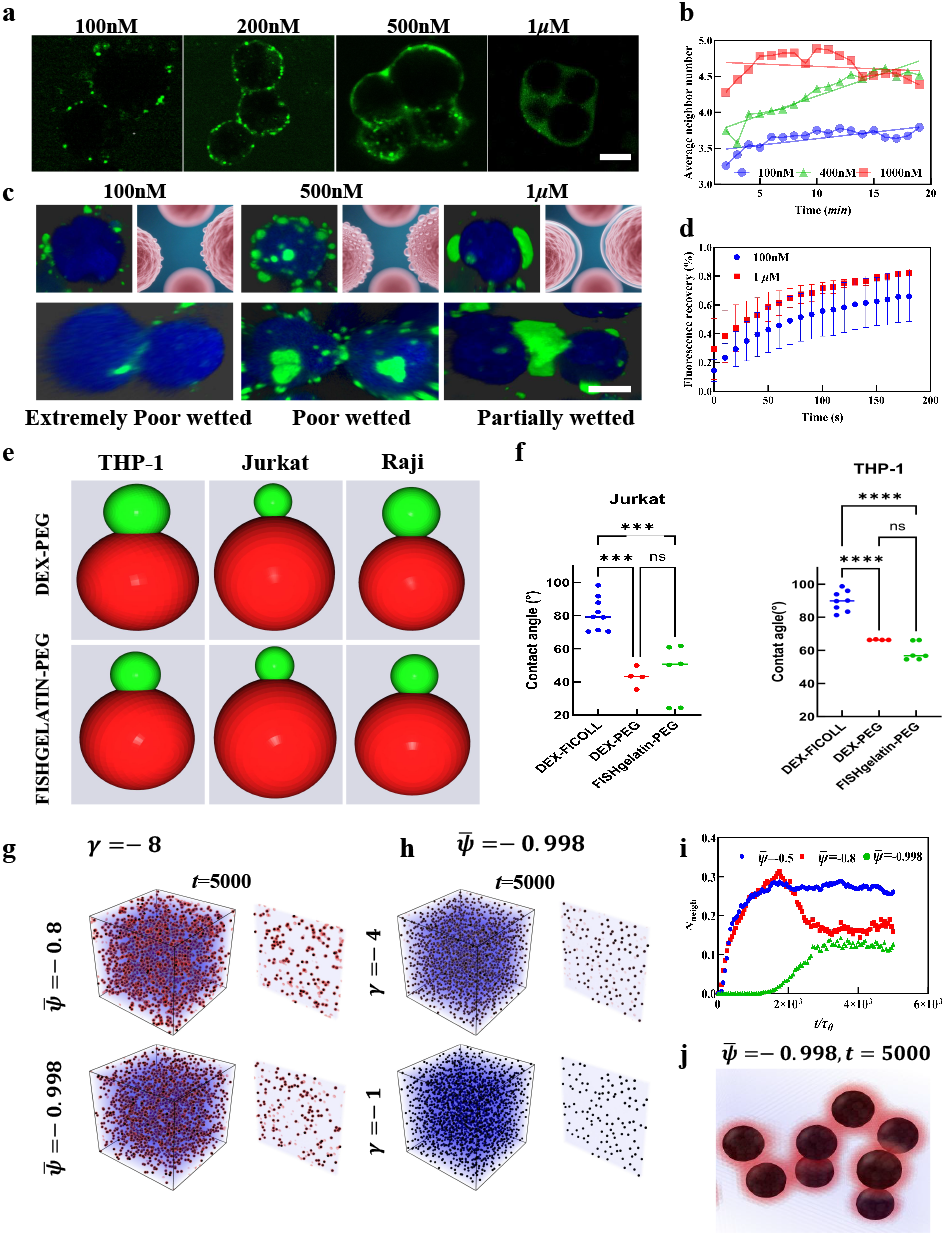
Wetting-induced aggregation is tunable by surface affinity and operates at nanomolar concentrations. **a** Confocal microscopy overview showing that FITC-tagged DEX (green) mediates cell aggregation in a concentration-dependent manner (100 nM to 1 *µ*M). Scale bar, 10 *µ*m. **b** Aggregation kinetics quantified as the average number of neighbors per cell over time for the conditions in **a. c** 3D Z-stack reconstructions of DEX condensates (green) on cell surfaces (blue). Scale bar, 10 *µ*m. **d** Fluorescence Recovery After Photobleaching (FRAP) analysis of the condensates. **e** 3D renderings of contact angles (*θ*) of the minority phase (green) on immune cell surfaces (red) in different ATPS systems (DEX–PEG and FISHGelatin–PEG). **f** Quantitative analysis of contact angles measured in **e**. Error bars represent mean ± s.d. (\*\*\*\**P <* 0.0001; ns, not significant; two-tailed *t*-test). **g** Simulations showing that strong wetting affinity (*γ* = −8) drives particle aggregation in the extreme case 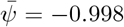 (*ϕ*_Cell_ = 2.5%). **h** Control simulations corresponding to **g**, showing that weak affinity (*γ* = −1 and −4) fails to induce aggregation. **i** Simulation kinetics: average neighbor number *N*_neigh_ as a function of time. **j** Simulation snapshot of capillary bridges between particles at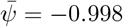 (*t* = 5000*τ*_0_).

Confocal microscopy further confirmed that at 100 nM, DEX appeared as punctate structures on the cell surface, consistent with extremely poor wetting (Fig. 2c). Strikingly, Fluorescence Recovery After Photobleaching (FRAP) analysis demonstrated that even under these dilute conditions, the condensates retained liquid-like properties, exhibiting substantial fluorescence recovery of ≈ 40% − 50% (Fig. 2d).

The aggregation observed on cell surfaces in the bulk one-phase regime reflects a wetting process, fundamentally governed by each cell’s surface affinity for the coexisting liquid phases. This affinity can be directly quantified by measuring the contact angle, *θ*, formed by a cell at the liquid–liquid interface [41–44]. To probe this key parameter and its role in aggregation, we developed an image analysis pipeline to precisely measure contact angles for multiple immune cell lines across different ATPS compositions (Fig. 2e–f; **Extended Data Fig**. 2a–b).

Quantitative analysis first confirmed the principle in our primary system. Jurkat cells displayed strong preferential wetting by the DEX phase in the canonical DEX–PEG system, with a low contact angle of *θ* ≈ 40° (Fig. 2f). To validate that this wetting preference drives aggregation, we examined Jurkat cells in two additional ATPS with distinct interfacial properties. In the DEX–FICOLL system, Jurkat cells exhibited poor wetting (*θ* ≈ 80°), which, as predicted by our framework, eliminated the attractive force and produced negligible aggregation (**Extended Data Fig**. 2d). By contrast, in the FISHgelatin–PEG system, Jurkat cells showed moderate wetting (*θ* ≈ 50°), and this composition was indeed able to support aggregation, even at bulk one-phase concentrations (**Extended Data Fig**. 2e and **Extended Data Fig**. 1c–d).

Importantly, surface affinity was not universal but cell-type specific. THP-1 cells exhibited their strongest preferential wetting in the FISHgelatin–PEG system (*θ* ≈ 55–65°), while showing only moderate affinity in the DEX–PEG system (*θ* ≈ 63–66°). Raji cells displayed a distinct “wetting fingerprint,” with high affinity for both the DEX–PEG (*θ* ≈ 45–50°) and FISHgelatin–PEG (*θ* ≈ 40–60°) systems (**Extended Data Fig**. 2c).

Collectively, these results demonstrate that immune cells possess distinct and tunable surface affinities, providing a quantitative physical basis for cell-specific aggregation and the competitive wetting phenomena explored later.

To elucidate the principles governing cellular self-organisation in a dilute extracellular milieu, we performed three-dimensional (3D) simulations of cell particles. We first examined how the volume fraction of the wetting phase, tuned by the average composition 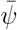, affects aggregate morphology under strong wetting affinity conditions (*γ* = −8). Remarkably, even when the wetting phase was extremely scarce and the system resided in the bulk one-phase regime 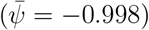, particles still underwent robust aggregation, forming structures comparable to those observed under less dilute, bulk binodal conditions (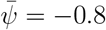; Fig. 2g). Quantitative analysis confirmed this similarity: the average neighbor number *N*_neigh_ was nearly identical in the one-phase and binodal regimes (Fig. 2i).

Crucially, aggregation in this dilute regime depended strongly on the wetting affinity of the minority phase. When the affinity was reduced (*γ* = −4 or *γ* = −1), particles remained dispersed at 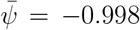 and failed to form aggregates (Fig. 2h). Detailed inspection of the aggregated microstructure revealed that the minority phase formed localised capillary bridges between particles at 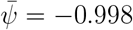 (Fig. 2j), mirroring our experimental observations.

Taken together, our experiments and simulations support the principles of heterogeneous nucleation. In the dilute minority-phase regime, the free-energy barrier (Δ*G*^∗^) for bulk phase separation is prohibitive. The cell surface, however, acts as a catalytic substrate that dramatically lowers this barrier. The reduction arises from the energetic gain of wetting: for surfaces with high affinity, the wetting energy (*f*_*s*_ ∝ *γψ*) substantially offsets the cost of creating a new interface. As a result, liquid-like condensates nucleate preferentially on cell surfaces, even under bulk one-phase conditions (Video 6). This surface-assisted nucleation mechanism explains the persistence of aggregation far below the bulk phase separation threshold and provides direct experimental validation of membrane wetting theories, which predict that confining phase separation to two-dimensional surfaces drastically reduces the saturation threshold required for nucleation [45, 46].

### Competitive Wetting Enables High-Fidelity Cell Recognition

Our finding that different cells exhibit distinct and tunable surface affinities (Fig. 2f) raises a critical question: can these physical differences be exploited to achieve specific intercellular recognition? To address this, we co-cultured cell lines with contrasting affinities (Jurkat–THP-1 and Jurkat–Raji) in the DEX/PEG ATPS. Strikingly, we found that recognition is exquisitely tuned by concentration. At high DEX concentration (1 *µ*M), the abundant wetting phase coated all cells non-selectively. By contrast, at an optimal low concentration (100 nM, bulk one-phase regime), high-fidelity recognition emerged: high-affinity cells formed homotypic clusters while excluding low-affinity neighbors (Fig. 3a).

**FIG. 3.**
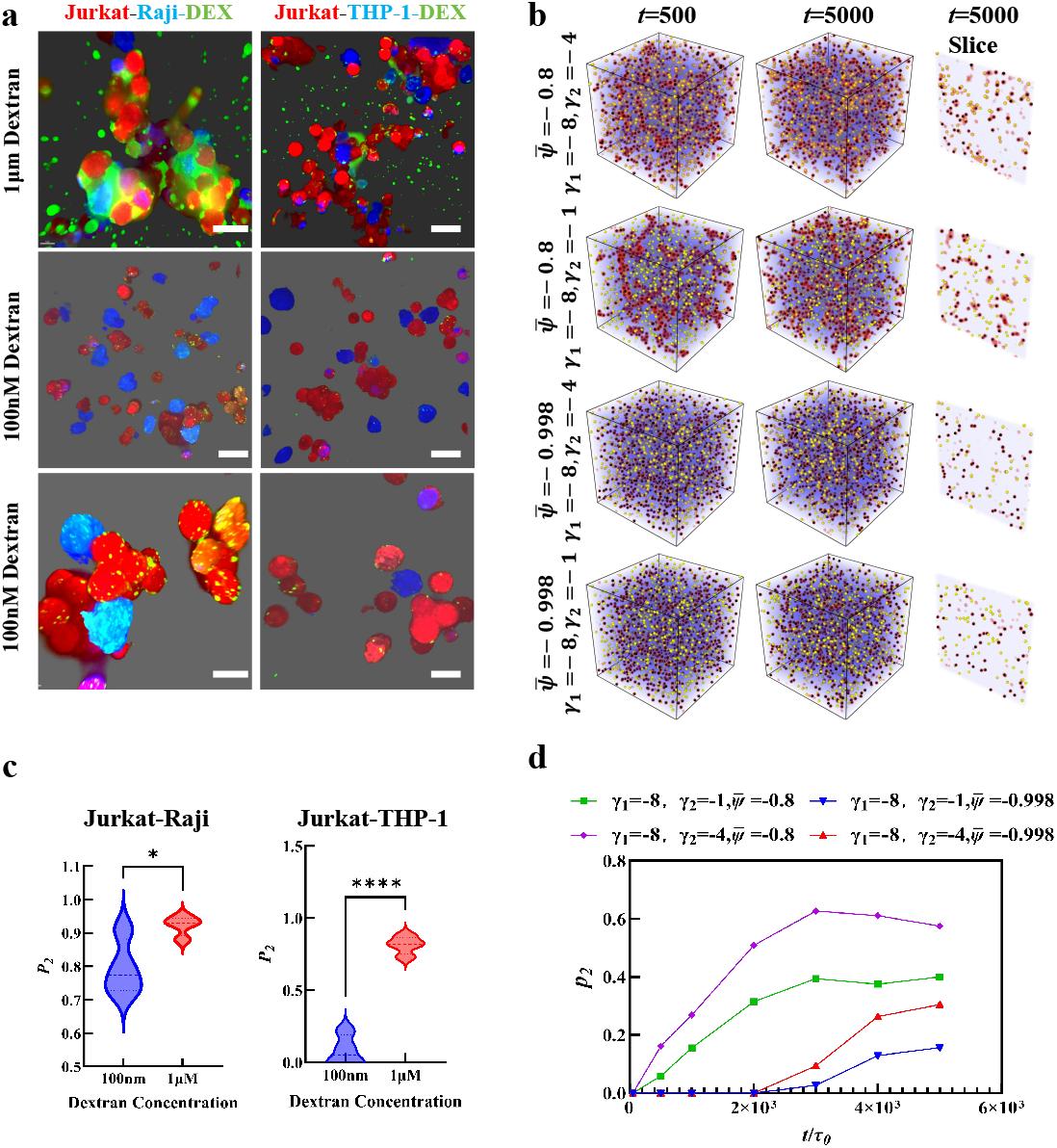
Competitive wetting drives specific intercellular recognition. **a** Confocal images of Jurkat–Raji and Jurkat–THP-1 co-cultures at high (1 *µ*M) and low (100 nM) DEX concentrations. Scale bar, 10 *µ*m. **b** 3D simulation snapshots of a binary particle mixture (*γ*_1_ = −8 versus *γ*_2_ = −4, −1) in phase-abundant 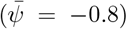 and resource-limited 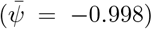 regimes. **C** Experimental quantification of sorting fidelity for Jurkat–Raji (left) and Jurkat–THP-1 (right) co-cultures. The sorting error rate *P*_2_ (fraction of low-affinity cells incorporated into clusters) is plotted for high (1 *µ*M) and low (100 nM) DEX concentrations. Lower *P*_2_ indicates higher specificity. \*\*\*\**P <* 0.0001, \**P <* 0.05. **d** Simulation-based analysis of sorting fidelity, showing *P*_2_ (fraction of low-affinity particles incorporated into aggregates) as a function of time *t*.

We hypothesised that this specificity arises from the principle of competitive wetting. To test this, we performed 3D simulations of binary mixtures comprising particles with high (*γ*_1_ = −8) and low (*γ*_2_ = −4, *γ*_2_ = −1) surface affinity for the wetting phase. The simulations revealed two regimes that mirror our experimental results (Fig. 3b). Under phase-abundant conditions 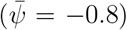, the system could afford the energetic cost of wetting both particle types, producing non-specific, heterotypic aggregates. In sharp contrast, under resource-limited one-phase conditions 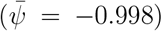, the scarce wetting phase was preferentially allocated to high-affinity particles, yielding exclusive homotypic aggregation.

This prediction was validated by quantitative analysis of sorting fidelity, *P*_2_, defined as the fraction of low-affinity cells incorporated into clusters. Both experiments (Fig. 3c) and simulations (Fig. 3d) demonstrated that the sorting error rate (*P*_2_) was dramatically reduced in the low-concentration, resource-limited regime, providing direct evidence for competitive wetting as the mechanism of high-fidelity recognition.

This behaviour can be rationalised by the principle of *competitive wetting*. To minimise free energy, scarce phase-separating molecules nucleate preferentially on the surface with the highest energetic payoff. Microscopically, this preference is determined by the wetting parameter *γ*, and macroscopically it manifests as a smaller contact angle *θ* in accordance with Young’s equation:

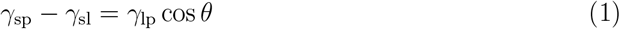

where *γ*_sp_ and *γ*_sl_ are the interfacial tensions between the cell surface and the sparse and dense liquid phases, respectively, and *γ*_lp_ is the tension between the two liquid phases. This “winner-takes-all” mechanism provides a physical basis for cell sorting, biasing a continuous spectrum of molecular affinities toward a digital-like “aggregate/reject” outcome. Thus, biological systems can exploit competitive wetting to perform sophisticated cellular screening simply by tuning the local extracellular concentration around a threshold.

### Interaction Range and Force Quantification

To understand the microscopic features of wetting-induced interactions, we quantified both the effective range and the magnitude of the forces involved. We first estimated the interaction range using serial dilution experiments, followed by direct force measurements with optical tweezers. In the dilution experiments, the cell volume fraction was converted into the average intercellular distance. Aggregation became negligible once the mean separation exceeded approximately five cell diameters (~50 *µ*m), providing an initial estimate of the effective range (Fig. 4a).

**FIG. 4.**
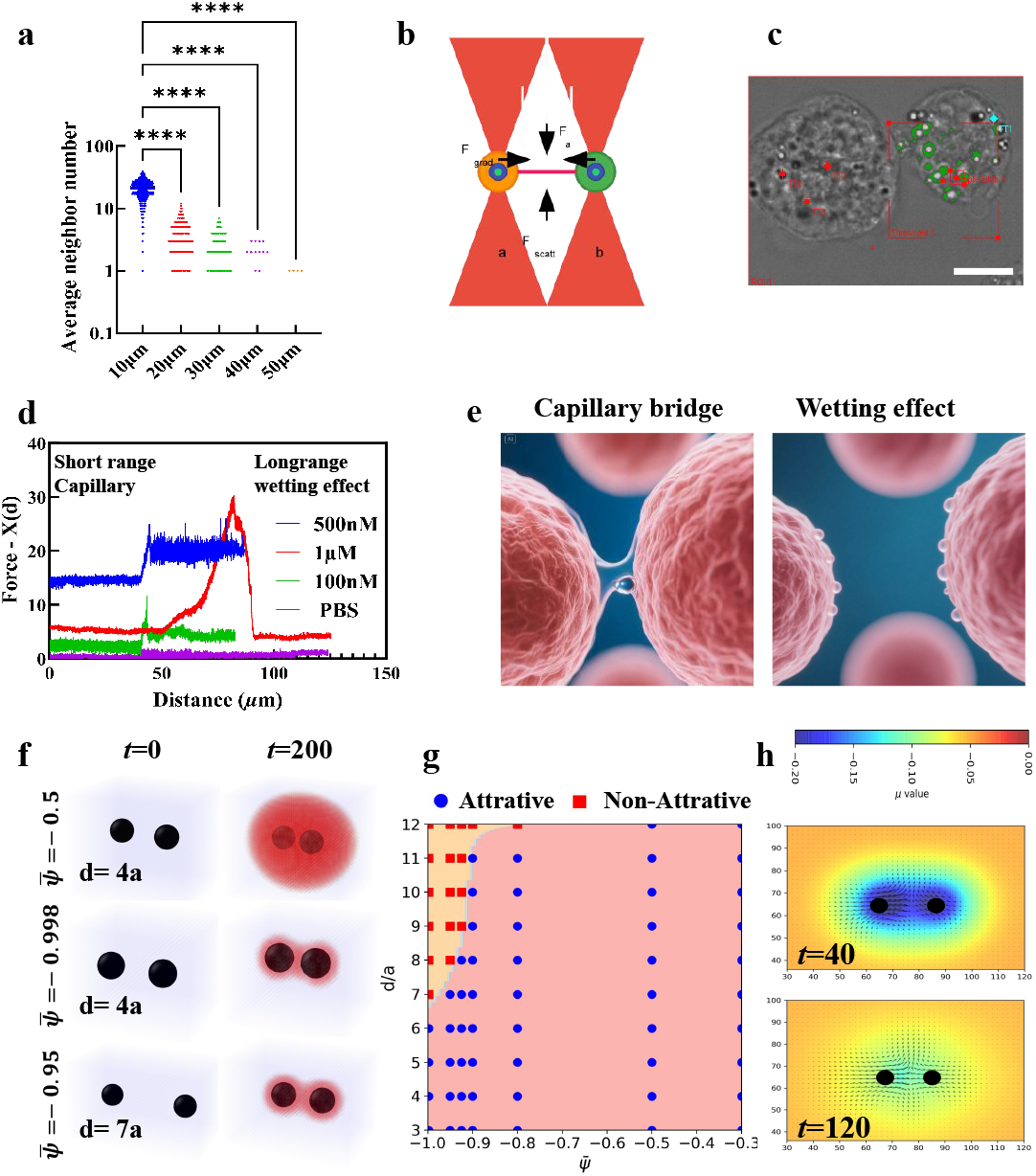
The interaction range and fundamental origin of wetting-induced adhesion force. **a** Experimental determination of the attractive force range. The plot shows the average number of neighbors as a function of the average intercellular distance, controlled by serial dilution of the cell suspension. **b** Schematic of the dual-trap optical tweezer setup used for direct force measurements between two cells (a and b). **c** Micrograph of two Jurkat cells bridged by a DEX condensate during a force measurement. Green circles indicate tracking traps; red circles indicate manipulation traps. Scale bar, 10 *µ*m. **d** Force–distance curves measured by optical tweezers at different DEX concentrations: 1 *µ*M and 500 nM (bulk two-phase regime), 100 nM (bulk one-phase regime), and PBS control. **e** Schematic illustrating particle aggregation near walls with affinity (*γ*_wall_ = −4). **f** Chemical potential field around two free particles in phase-abundant 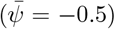 and resource-limited (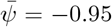, −0.998) regimes with different initial separations *d*. **g** Simulated state diagram mapping interaction states (attractive vs. non-attractive) as a function of normalised particle separation (*d/a*) and average composition 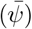 **h** Simulation snapshots showing the temporal evolution of the chemical potential around two particles at 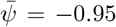 at early (*t* = 40) and late (*t* = 120) times. 21

We then employed a dual-trap optical tweezer setup to directly measure cell–cell adhesion in the phase-separating medium (Fig. 4b–c). By pulling apart two aggregated cells, we recorded force–distance curves that revealed attractive forces on the order of piconewtons (*p*N) (Fig. 4d). At 1 *µ*M DEX, the profile displayed two distinct components: a strong, short-range ‘capillary’ force peaking when the liquid bridge was stretched, and a sustained, long-range ‘wetting-effect’ force that plateaued over distances up to 80 *µ*m (Fig. 4e). Correcting for initial cell overlap, this corresponds to an effective interaction range of about six cell diameters, consistent with our dilution-based estimate. At lower concentrations (100 and 500 nM, bulk one-phase regime), the same two-component signature was observed, though both magnitude and range were reduced, with interactions limited to ~50 *µ*m. As expected, no adhesive force was detected in PBS controls.

To generalise these findings, we performed 3D simulations of two free particles in a phase-separating mixture. The results revealed that the interaction range is a highly nonlinear function of the average composition 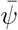, which dictates the available wetting-phase volume (Fig. 4f–g). In the “resource-limited” regime 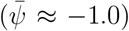, scarce wetting phase permitted only localised bridges at close approach, yielding short-range interactions. In contrast, under “phase-abundant” conditions 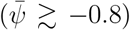, extensive wetting layers formed, enabling long-range capture of particles separated by more than five radii. This principle — that the capture range scales with phase availability — aligns with our experimental observation that larger, denser aggregates retain more neighbors (Fig. 4a).

Finally, to identify the origin of this long-range attraction, we examined the evolution of the chemical potential field *µ* (Fig. 4h). Each cell establishes an inhomogeneous *µ* field due to surface affinity, generating a gradient (∇*µ*) that drives inward flux of the wetting phase. This gradient induces not only diffusional material transport (*J* = −*L*∇*µ*) but also hydrodynamic flow. When two cells approach, reduced flux into the intercellular gap creates a spatial imbalance, producing a net osmotic pressure that pulls cells together —a process we describe here as wetting-induced attraction, analogous in spirit to capillary or Casimir-like forces in soft matter systems [4, 5, 38, 47–49]. The resulting flow brings cells into close contact, where a stable, lower-energy capillary bridge can form as the *µ* field equilibrates.

In summary, wetting-induced interactions operate via two distance-dependent regimes. At long range, cells experience attraction mediated by chemical-potential gradients and wetting flows, extending over more than five diameters. At short range, direct capillary bridges generate strong adhesive forces. The magnitude and range of both regimes are governed by the volume of the wetting phase and its affinity for cell surfaces.

### Adhesion of Suspended Cells to Surfaces and the Role of Shear Flow

Having established that competitive wetting governs cell–cell interactions, we next examined a biologically critical scenario: the adhesion of suspended cells to an adherent surface, a process relevant to leukocyte extravasation. We hypothesised that surface geometry itself introduces a critical asymmetry that dictates adhesion efficiency. To test this, we first used a non-adherent petri dish with suspended cells in DEX/PEG mixtures of varying concentrations. In the absence of phase-separating agents, cells remained suspended (**Extended Data Fig**. 3a), confirming the lack of intrinsic attraction to the surface. At low DEX concentration (100 nM in 5 wt.% PEG), however, cells adhered strongly to the wall (**Extended Data Fig**. 3b). In contrast, when DEX was increased to 1 *µ*M, above the spinodal threshold, cells detached and became suspended within the bulk-separated liquid (**Extended Data Fig**. 3c). We then validated this effect in a physiological context using co-cultures of suspended Jurkat cells and an adherent layer of human cardiac microvascular endothelial cells (HCMECs), which was tested with high affinity with DEX phase (Data not shown). Consistent with our earlier observations, Jurkat cells were efficiently recruited to the HCMEC layer at low DEX concentrations (100 nM, one-phase regime; **Extended Data Fig**. 4a). Recruitment was mediated by puncta-like DEX condensates on the surface (**Extended Data Fig**. 3d–e) and diminished when the wetting phase was abundant.

Our quasi-2D simulations reproduced these phenomena. At phase-abundant composition 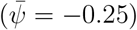, suspended particles near a neutral flat substrate remained largely dispersed in the bulk fluid. By contrast, under both bulk binodal and one-phase conditions (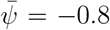 and −0.998), particles were rapidly captured onto the wall, forming dense surface-bound aggregates (**Extended Data Fig**. 4b; Fig. S11). Kinetic analysis revealed sharp decreases in the average particle–wall distance *d*_wall_ (**Extended Data Fig**. 4d), accompanied by rapid increases in the average neighbor number *N*_neigh_ (**Extended Data Fig**. 4c). Thus, in resource-limited regimes 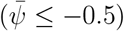, the wall acts as a strong attractor, pulling particles from the bulk and facilitating their subsequent aggregation. The underlying mechanism arises from spatially inhomogeneous chemical potential fields (**Extended Data Fig**. 4f, 4g): lower *µ* near particle and wall surfaces compared to the bulk establishes gradients (∇*µ*) that drive both diffusive flux and hydrodynamic flow, transporting the minority phase toward surfaces.

We next investigated how wall and particle affinities modulate adhesion. Adhesion slowed when particle–phase wetting affinity was weaker (**Extended Data Fig**. 5). By contrast, introducing a wall with preferential affinity for the wetting phase (*γ*_wall_ = −4) greatly accelerated adhesion (**Extended Data Fig**. 4e, 4h, **and** **Extended Data Fig**. 6). Under these conditions, the wetting phase coated the wall (**Extended Data Fig**. 4e), creating a thick layer that captured particles efficiently. Quantitative analysis confirmed that attractive walls (*γ*_wall_ = −4) produced much faster accumulation compared with neutral walls (**Extended Data Fig**. 4h).

In biological settings, cells are often subjected to external flow fields and shear forces, which can significantly modulate their aggregation behaviour. As depicted in **Extended Data Fig**. 4i, a striking observation was that the presence of an external shear force dramatically accelerated the kinetics of particle aggregation. For instance, comparing *f*_*d*_ = 0 (no shear) with *f*_*d*_ = 5 × 10^−5^ at the same condition of 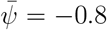 (first two rows of **Extended Data Fig**. 5), the system with shear rapidly transitioned from a dispersed state to a fully aggregated state within *t* = 2000*τ*_0_, whereas the no-shear condition required a longer time to achieve similar levels of aggregation. Similar accelerations were observed with different shear magnitudes and interaction strengths. This acceleration can be attributed to increased collision frequencies between particles under shear, as well as shear-induced particle-phase mixing which facilitates the formation of a wetting layer and subsequent bridging.

Together, these findings demonstrate that wetting-induced adhesion to surfaces is robust, tunable by both phase availability and interfacial affinity, and effective even under physiological shear (Video 7). This provides a physical basis for the rapid arrest of suspended cells on endothelium and other surfaces in vivo [50–53].

### Biological Function: Phase Separation as a Physical Regime for Cell Aggregation

Our findings thus far establish a robust physical framework for cell adhesion driven by wetting and capillary forces. A critical test is whether endogenous biomolecules employ this mechanism to generate functional consequences in biological contexts. To address this, we examined proteins known to undergo extracellular LLPS and mediate cell communication — Galectin-3 [23, 54, 55] and the chemokine CCL5 [19] — and assessed their ability to drive cell aggregation.

Confocal imaging revealed a clear concentration dependence for Galectin-3 (GAL3) in mediating cell clustering. It has previously been established that GAL3 can form clusters on the surface of T cells [23]. Consistent with this, as the concentration of GFP-tagged GAL3 was reduced from 10 *µ*M (Fig. 5a, 5b) to 1 *µ*M (Fig. 5c, Video 8) and then to 100 nM (Fig. 5d), the extent of cell clustering decreased progressively. At 10 *µ*M, cells formed large, compact clusters, their surfaces extensively coated with bright green condensates. High-resolution 3D reconstructions confirmed that these condensates were enriched at cell–cell junctions, effectively acting as “liquid glue” (Fig. 5b). Control experiments validated the specificity of this phenomenon (Fig. 5e). Robust clustering occurred at 1 *µ*M and 10 *µ*M GAL3, with weaker aggregation at 0.1 *µ*M. By contrast, PBS buffer or a phase-separation– deficient GAL3 mutant failed to induce clustering, confirming that the effect is mediated directly by LLPS (Video 9).

**FIG. 5.**
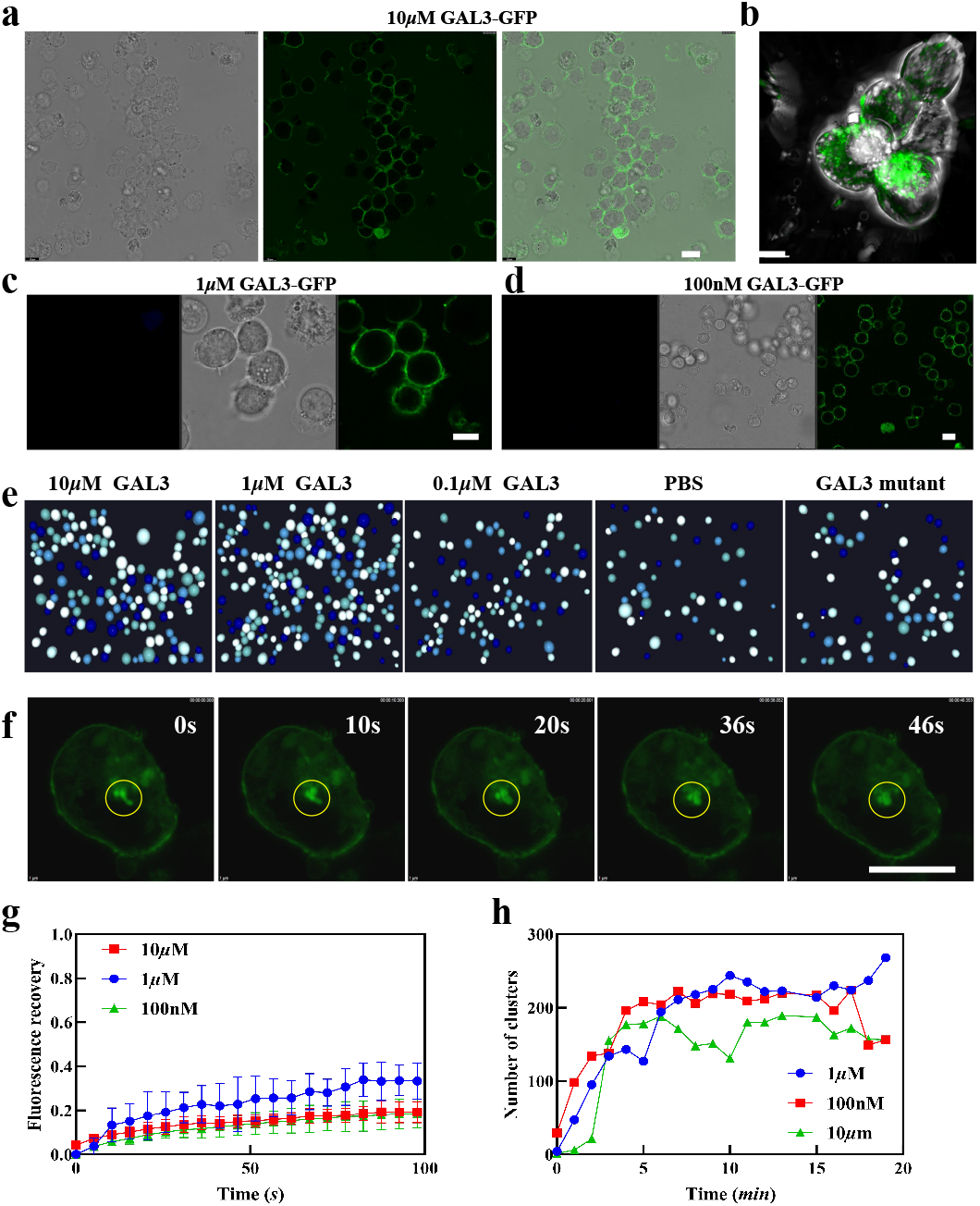
Endogenous phase-separating proteins mediate functional cell aggregation via the wetting-induced attraction mechanism. **a** Bright-field and confocal images of Jurkat cells incubated with a high concentration (10 *µ*M) of GFP-tagged Galectin-3 (GAL3-GFP), showing extensive cell aggregation and surface coating. Scale bar, 10 *µ*m. **b** High-resolution 3D reconstruction of a cell cluster from **a**, revealing GAL3-GFP condensates (green) at cell–cell junctions, acting as a “liquid glue”. Scale bar, 5 *µ*m. **c** Aggregation and surface wetting observed at an intermediate GAL3-GFP concentration (1 *µ*M). Scale bar, 10 *µ*m. **d** Aggregation and surface wetting observed at a low GAL3-GFP concentration (100 nM). Scale bar, 10 *µ*m. **e** Validation of LLPS-dependent aggregation. Robust clustering is observed at 10 *µ*M and 1 *µ*M GAL3, but aggregation is abrogated in PBS buffer and with an LLPS-deficient GAL3 mutant. **f** Time-lapse microscopy showing the liquid-like behaviour of condensates: two smaller droplets coalesce into a larger spherical droplet over ~46 s. **g** Quantitative FRAP curves of GAL3-GFP on Jurkat cell surfaces at different concentrations. **h** Aggregation kinetics, tracking the number of clusters over time and showing that higher GAL3 concentrations lead to faster and more extensive aggregation.

We characterised the physical nature of these condensates. Time-lapse microscopy showed two smaller GAL3 condensates fusing into a single droplet within ~46 seconds — a hallmark of liquid-like behaviour (Fig. 5f). Fluorescence Recovery After Photobleaching (FRAP) demonstrated substantial recovery (*>*40% within 100 seconds) across all concentrations, indicating high internal mobility (Fig. 5g; **Extended Data Fig**. 7a). Kinetic analysis further showed that aggregation rates scaled with protein concentration, with higher GAL3 accelerating cluster formation (Fig. 5h).

To demonstrate generality beyond GAL3, we also examined CCL5. As previously reported [19], CCL5 aggregates cells only when presented at an appropriate molar ratio with heparin sulfate (*>*1:5). At 2 *µ*M and 500 nM, CCL5 indeed drove Jurkat aggregation in the presence of heparin sulfate, producing small condensates (**Extended Data Fig**. 7c).

Together, these results establish that endogenous biomolecules employ wetting-induced aggregation as a physical mechanism to mediate cell–cell interactions. Galectin-3 and CCL5 exemplify how extracellular LLPS can function in a specific, concentration-dependent manner to promote multicellular organisation.

### LLPS-Mediated Bridges Accelerate Contact-Dependent Signaling

We next asked whether LLPS-mediated aggregation accelerates biological processes that require stable cell–cell contact. To this end, we used a Jurkat–Raji co-culture engineered with a CD27/CD70 contact-dependent signaling reporter. In this system, one cell line presents the ligand (CD70) and the other the receptor (CD27), with their interaction producing a quantifiable luminescent signal (measured in relative luminescence units, RLU). This assay provides a direct readout of intercellular presentation efficiency. We hypothesised that LLPS-generated bridges act not only as physical tethers but also as dynamic synapses that facilitate and accelerate signaling (Fig. 6a, 6b).

**FIG. 6.**
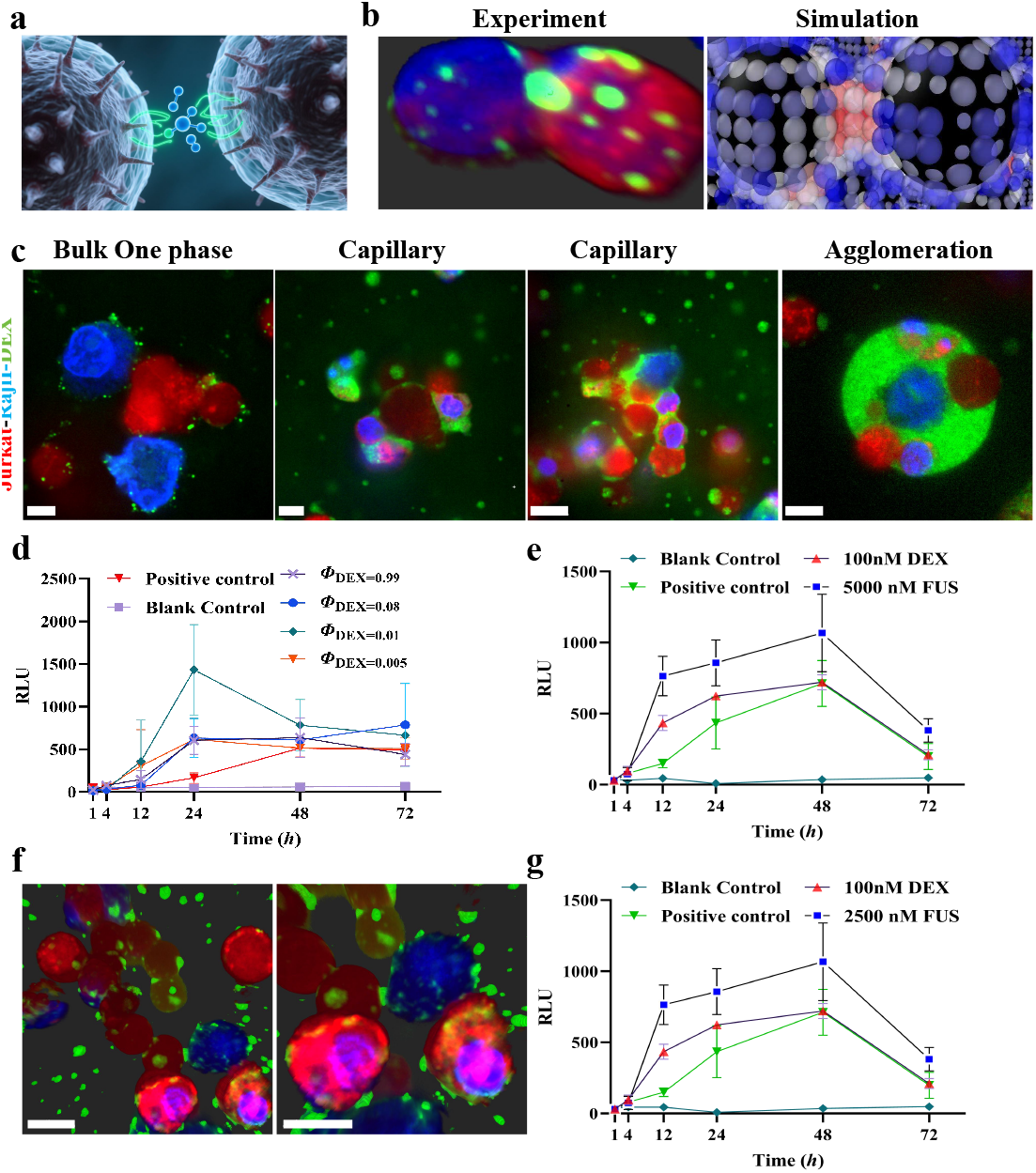
LLPS-mediated bridges as physical catalysts for intercellular signaling. **a** Conceptual illustration of biomolecular condensate-mediated cell–cell contact and signaling enhancement, using a CD27/CD70 Jurkat–Raji reporter system to demonstrate that LLPS-driven bridges enhance intercellular signaling efficiency. **b** Comparison of experimental and simulated cell–cell interactions mediated by a phase-separating medium. **c** Confocal images of the reporter system mediated by DEX (green, DEX–FITC). Jurkat cells (red, Cytotell Orange) and Raji cells (blue, Cytotell Blue) interact in different morphological regimes. Scale bar, 10 *µ*m. **d** Reporter signal kinetics (RLU vs. time) in the bulk binodal regime. Various DEX volume fractions (*ϕ*_DEX_) all show significant signal enhancement compared to blank and positive controls. **e** Reporter signal kinetics in the bulk one-phase regime. Nanomolar DEX concentrations (200 nM, 400 nM) induce rapid activation, with initial kinetics even faster than those in the binodal regime. **f** Confocal microscopy images of Jurkat (red) and Raji (blue) cells in the bulk one-phase regime mediated by FUS protein. Scale bars, 10 *µ*m. **g** Quantification of downstream signaling (CD27/CD70 Jurkat–Raji reporter assay, RLU) in a co-culture system with 2500 nM FUS protein over time, compared to positive and blank controls and to higher DEX concentration (100 nM).

We first tested this under bulk two-phase conditions (Fig. 6c), corresponding to the morphologies described in Fig. 1a. All phase-separating compositions significantly accelerated and amplified reporter activation relative to the blank control (Fig. 6d). Compositions yielding capillary states (*ϕ*_DEX_ = 0.01, 0.005) produced especially rapid and robust signals, confirming that LLPS-mediated aggregation strongly promotes contact-dependent signaling.

Strikingly, the effect persisted even in the bulk one-phase regime. At nanomolar DEX concentrations (200–400 nM), where only localised bridges were expected, the reporter signal remained robustly activated (Fig. 6e). Although the maximum intensity was slightly lower than in binodal conditions, the initial activation rate was faster. We attribute this to the resource-limited regime: the scarcity of wetting phase forces cells into more intimate contacts (Fig. 6c, “bulk one phase”), raising the effective local concentration of CD27/CD70 pairs and accelerating the initial kinetics.

Finally, to test whether this catalytic principle extends to endogenous proteins, we examined FUS, an RNA-binding protein known to undergo LLPS, which would not interfere with the extracellular biological function of the cell. Confocal microscopy confirmed that FUS could also form condensates bridging Jurkat and Raji cells in a capillary-like manner (Fig. 6f). Remarkably, at 2.5 *µ*M, FUS induced rapid, robust reporter activation comparable in both speed and magnitude to the optimal DEX condition (Fig. 6g).

These results demonstrate that the acceleration of intercellular signaling is a generalizable consequence of extracellular phase separation, independent of the molecular identity of the condensate. Any biomolecule capable of undergoing LLPS can, in principle, act as a physical catalyst to promote functional cell–cell interactions.

## SUMMARY AND OUTLOOK

In this study, we introduce and validate a new physical framework for the earliest stages of intercellular adhesion, moving beyond the classical receptor–ligand paradigm. We demonstrate that extracellular LLPS transforms the pericellular environment into a dynamic three-phase system, generating powerful non-equilibrium forces that drive rapid, long-range, and selective cell–cell interactions. This biophysical perspective provides compelling answers to long-standing questions of how non-motile cells achieve rapid targeting and capture in crowded environments such as the bloodstream.

Our results reveal that wetting is a key principle governing multicellular organisation within a phase-separating milieu. By bridging concepts from soft-matter physics and cellular biology, we uncover a mechanism by which cells transition from sparse individual states to highly interconnected collectives. Notably, we identify a potent adhesion mechanism that remains active at biologically relevant nanomolar concentrations, where the system resides in a bulk one-phase regime. In this regime, cell surfaces with strong affinity for a phase-separating component act as nucleation templates, generating wetting-induced attraction. This force arises from chemical-potential gradients that drive hydrodynamic flow. It bears similarity to Casimir forces in binary mixtures [30, 56, 57], yet unlike the fluctuation-limited Casimir case, it extends over many cell diameters and thereby enables rapid cell capture. This non-equilibrium dynamics may help explain how cells can capture neighbors over distances up to ~60 *µ*m — far beyond the nanometre-scale reach of conventional receptor–ligand interactions [4, 5, 38, 47–49].

This framework also provides a new physical basis for biological specificity through the principle of competitive wetting. Unlike the “lock-and-key” recognition of receptor–ligand binding, competitive wetting is a thermodynamic selection process [2, 3]. Scarce phase-separating molecules preferentially condense on the surface with the highest energetic pay-off — defined by strong affinity (*γ*) and low contact angle (*θ*). This “winner-takes-all” mechanism effectively translates a continuous spectrum of affinities into a digital-like aggregate/reject outcome, enabling biological systems to toggle interactions from promiscuous to highly specific simply by modulating local molecular concentrations.

Beyond cell–cell adhesion, our model redefines cell–surface interactions through curvature-dependent nucleation. Curvature alters the balance between interfacial energy cost and wetting gain: on highly curved cell surfaces the nucleation barrier is high, whereas on low-curvature substrates it is reduced. As a result, phase separation preferentially wets flat surfaces such as endothelial walls, which then recruit nearby cells via capillary bridges. This offers a purely physical explanation for the rapid capture of circulating cells during processes such as leukocyte extravasation or cancer metastasis, potentially preceding or acting in parallel with selectin–integrin cascades [58, 59]. Crucially, we demonstrate that endogenous proteins including Galectin-3 and CCL5 can harness this mechanism, firmly situating the framework within a physiological context.

Functionally, LLPS-driven aggregation acts as a biological catalyst. First, by overcoming the spatial barrier, it dramatically increases the frequency of productive cell encounters. Second, by creating condensates at cell–cell junctions, it overcomes the concentration barrier, enriching ligands and receptors at the contact site. This dual catalytic role — kinetic and chemical — illustrates how cells exploit non-equilibrium physics to orchestrate rapid and robust functional outcomes.

In conclusion, our work suggests that cell adhesion is not solely a biochemical event but the synergistic outcome of molecular recognition and emergent biophysical forces. By considering the cell–environment interface as a dynamic three-phase system, we provide a complementary physical model that helps account for the speed, range, and specificity of initial adhesion. This perspective not only reshapes our understanding of multicellular organisation but also opens new therapeutic avenues: by modulating the phase behaviour of the extracellular environment, it may be possible to control cell–cell interactions in immunity, development, and disease.

## METHODS

### Cell Lines and Culture Conditions

Human monocyte cell line THP-1, Raji, and Jurkat cells were obtained from Procell. Raji and Jurkat cells were routinely maintained in RPMI-1640 medium supplemented with 10% (v/v) fetal bovine serum and 1% (v/v) penicillin-streptomycin. THP-1 cells were cultured in RPMI-1640 medium supplemented with 10% (v/v) fetal bovine serum, 0.05 mM 2-Mercaptoethanol, and 1% (v/v) penicillin-streptomycin. Human T-lymphocyte cell line Jurkat (CD27 reporter, InvivoGen, Cat #jktl-cd27) and B-lymphocyte cell line Raji (CD70 ligand, included in InvivoGen Cat #jktl-cd27) were cultured in Iscove’s Modified Dulbecco’s Medium (IMDM, InvivoGen) supplemented with 2 mM L-glutamine, 25 mM HEPES, and 10% (v/v) FBS.

Human Cardiac Microvascular Endothelial Cells (HCMEC) were cultured in Endothelial cell culture medium. Cells were passaged every 2-3 days using 0.25% Trypsin-EDTA and used at passages 4-6.

HEK293 cells were cultured in DMEM medium supplemented with 10% FBS and 1% P/S. All cells were maintained at 37°C in a humidified atmosphere containing 5% CO_2_. Cells were routinely tested for mycoplasma contamination and were confirmed negative.

### Preparation of Aqueous Two-Phase Systems (ATPS)

Aqueous two-phase systems were prepared using Dextran (DEX, MW = 500 kDa, and Polyethylene Glycol (PEG, MW = 100 kDa). Stock solutions of DEX and PEG were prepared at different concentrations in 1x Phosphate Buffered Saline (PBS) pH 7.4 (composition: 137 mM NaCl, 2.7 mM KCl, 10 mM Na_2_HPO_4_, 1.8 mM KH_2_PO_4_) or relevant cell culture medium.

For experiments mapping morphological phase space, stock solutions were mixed to achieve the desired final volume fractions (*ϕ*). The total volume of the system was defined as *V*_total_ = *V*_DEX_ + *V*_PEG_ + *V*_cells_. The volume fractions of dextran and cells were then calculated as *ϕ*_DEX_ = *V*_DEX_*/V*_total_ and *ϕ*_Cell_ = *V*_cells_*/V*_total_, respectively. The total volume of cells (*V*_cells_) was determined by first counting the number of cells using a hemocytometer, and then calculating the aggregate volume by idealizing each Jurkat cell as a sphere with a diameter of 10 *µ*m.

For experiments in the nanomolar regime, a 500 kDa DEX stock solution was prepared in relevant cell culture medium at 5 wt% concentration. This stock was then serially diluted with a 5 wt.% PEG solution (prepared in cell culture medium) to achieve final DEX concentrations ranging from 50 nM to 1000 nM. The binodal curve was determined by cloud point titration at various temperatures until visual turbidity appears.

Other ATPS systems, including DEX-FICOLL and FISHgelatin-PEG, were prepared similarly by mixing the polymers in PBS or cell culture mediums.

### Cell Aggregation Experiments

Cells were harvested by centrifugation (300 x g) for 5 min, washed twice with 1x PBS, and resuspended in the prepared ATPS solutions or solutions containing phase-separating proteins (Galectin-3, CCL5, FUS). The final cell volume fraction was typically *ϕ*_Cell_ = To allow proper dispersion of the phases, cell suspensions were gently pipetted before observation.

The cell suspension was loaded into handmade petri dishes with proper sealing to avoid evaporation. Aggregation kinetics were monitored using time-lapse microscopy (Leica DMi8) with a 40x air objective.

For competitive wetting experiments, two cell lines were pre-labeled with distinct fluorescent dyes (Cytotell Orange and Cytotell Blue, AAT Bioquest, Cat# 22270 & 22271) according to the manufacturer’s instructions. Labeled cells were mixed at a 1:1 ratio and resuspended in DEX/PEG solutions at 100 nM or 1 *µ*M DEX. Sorting fidelity (*P*_2_, the fraction of low-affinity cells incorporated into clusters) was quantified manually.

### Image Analysis of Cell Aggregation

Time-lapse bright-field or fluorescence images obtained from cell aggregation assays were quantitatively analyzed using a custom Python script to characterise aggregation kinetics and morphology. The analysis pipeline involved the following steps:

1. **Image Pre-processing:** Raw images were first converted to grayscale.
2. **Cell Segmentation:** Individual cells were identified using Cellpose (Version 3.0, with model: ‘cyto’) for robust individual cell segmentation.
3. **Cluster Identification:** Cell clusters were identified based on a distance-based criterion. Cells were considered part of the same cluster if the distance between their centroids was less than or equal to 1.5 times the average cell diameter (15 *µ*m). This was implemented using DBSCAN with specified parameters.
4. **Quantitative Aggregation Metrics:** For each image and time point, the script calculated:
  - Total Cell Count (*N*_total_): The total number of individually segmented cells.
  - Number of Clusters (*N*_clusters_): The total count of identified cell clusters.
  - Average Cluster Size (*S*_avg_): The mean number of cells per cluster, calculated as *N*_total_*/N*_clusters_.
  - Fraction of Aggregated Cells (*F*_agg_): The proportion of cells participating in clusters containing more than 3 cells, calculated as (Total cells in clusters - Number of clusters) / Total cells.
  - Cluster Size Distribution: The frequency distribution of clusters with different numbers of cells.

### Contact Angle Measurement

To quantify cell surface wettability, cells were introduced into a pre-equilibrated ATPS at a composition of 5 wt.% DEX, 5 wt.% PEG in PBS. Cells were allowed to sediment and adsorb to the liquid-liquid interface for 15 minutes at room temperature. Images of cells at the interface were acquired using a Leica Stellaris 8 confocal microscope with a 63x/1.4 NA oil immersion objective.

After imaging, Cellpose (Version 3.0) was used for automated segmentation of individual cells and the surrounding liquid-liquid interface. Default or optimised parameters were applied for cell body and interface detection. Second, the segmented boundaries were imported into OPEN3D (Version 0.15.1) to reconstruct the 3D geometry of the cell and the interface. From this 3D reconstruction, the precise three-phase contact line (where cell, DEX-rich phase, and PEG-rich phase meet) was identified. Finally, a custom Python script was employed. This script performed geometric fitting of the cell surface and the liquid-liquid interface near the contact line to calculate the local tangent vectors. The contact angle (*θ*) was then derived from the angle between these tangent vectors, consistent with Young’s equation. The angle *θ* is reported relative to the DEX-rich phase. At least 10 cells were analyzed for each condition.

### Optical Tweezer Force Measurement

A dual-trap optical tweezer system (Tweeze 305, OptoSigma) and an inverted microscope (Nikon Ti-E) equipped with an infrared laser (1064 nm, IPG Photonics, up to 10 W) and a 100x/1.3 NA water immersion objective was used to measure the interaction forces between cell pairs.

1. **Calibration:** The trap stiffness for each optical trap was calibrated using the power spectral density analysis of the Brownian motion of trapped cells. The typical trap stiffness used was 0.26 pN/nm.
2. **Experimental Procedure:** Jurkat cells were suspended in DEX/PEG solutions (100 nM, 500 nM, 1 *µ*M DEX, all in PBS) or PBS control. Two individual cells were captured by separate optical traps and brought into contact. After allowing the capillary bridge to stabilise for 10 minutes, the movable trap was retracted at a constant velocity of 1 *µ*m/s.
3. **Data Acquisition and Analysis:** The positions of the trapped cells were tracked using a quadrant photodetector (QPD, Thorlabs). The force exerted on the cell was calculated by multiplying the displacement of the cell from the trap center by the trap stiffness. Force-distance curves were generated, and the interaction range and magnitude of the ‘capillary’ and ‘wetting-effect’ forces were analyzed using Python.

### Cell-Surface Adhesion

For static adhesion assays, suspended Jurkat cells were introduced into non-adherent confocal petri dishes (Treated with anti-adhesion rinse solution) or onto a confluent monolayer of HCMECs cultured on confocal petri dishes. Cells were resuspended in DEX/PEG mixtures (100 nM to 5 *µ*M DEX, in PBS). Adhesion was assessed after 30 minutes by acquiring images using confocal microscopy.

### Protein Expression and Purification

#### GAL3

The wild-type human galectin-3 gene was cloned from cDNA. The LLPS-deficient WY/G mutant was synthesised based on a previous report [55]. Both constructs were cloned into a modified pET-32a vector with an N-terminal 6xHis-GFP fusion tag. Plasmids were transformed into *E. coli* BL21 (DE3) cells. Cells were grown in LB medium at 37°C to an OD_600_ of 0.6-0.8. Protein expression was induced with 0.5 mM IPTG at 18°C for 16 hours. Cells were harvested by centrifugation and lysed by sonication on ice in lysis buffer (50 mM Tris-HCl pH 7.5, 300 mM NaCl, 10 mM Imidazole, 5% Glycerol, 1 mM PMSF, 1 mM DTT). The lysate was clarified by centrifugation. The protein was purified using Ni-NTA affinity chromatography followed by size-exclusion chromatography on an AKTA system (Cytiva, Model). The final buffer composition was 20 mM Tris-HCl pH 7.5, 150 mM NaCl, 1 mM DTT, 5% Glycerol. Protein concentration was determined by Bradford assay and stored at −80°C.

#### FUS

The FUS expression plasmid (Addgene #183236) was used for transient transfection of HEK293 cells using Polyethylenimine. Cells were cultivated for 4-6 days post-transfection and lysed by sonication in lysis buffer (50 mM Tris-HCl pH 7.4, 150 mM NaCl, 1 mM PMSF, 1 mM DTT). The lysate was purified using Ni-NTA affinity chromatography. The purified protein was dialyzed into the final buffer (PBS with 10% glycerol, pH 7.4).

#### CCL5

The CCL5 protein was kindly provided by the Luo group, purified according to reported protocols [Reference for purification method].

### Confocal Microscopy and Fluorescence Recovery After Photobleaching (FRAP)

Imaging was performed using a confocal laser scanning microscope Leica Stellaris 8. Fluorescent markers were excited using 488 nm and detected through appropriate emission filters.

FRAP experiments were conducted to assess the fluidity of DEX (Fig. 2d) and GAL3-GFP condensates. A region of interest (ROI) within the condensate was bleached using the 488 nm laser at 10% power intensity (0.40 mW) for 10 seconds. Fluorescence recovery was monitored by time-lapse imaging at every 10s for 100s. The fluorescence intensity within the bleached ROI was normalised to the pre-bleach intensity and corrected for background photobleaching, then fitted to a single exponential recovery model to extract the mobile fraction and half-time of recovery.

### Contact-Dependent Signaling Assay (CD27/CD70 Reporter)

A Jurkat-Raji co-culture system engineered for CD27/CD70 contact-dependent signaling was utilised. Specifically, we used the hCD27-BIONF Reporter Cells (InvivoGen, Cat #hcd27-onf), which consist of:

- **Jurkat-CD27 Reporter Cells:** Genetically engineered Jurkat cells that stably express the human CD27 receptor and a NF-*κ*B-inducible secreted embryonic alkaline phosphatase (SEAP) reporter gene. Upon engagement of CD27 by its ligand CD70, NF-*κ*B signaling is activated, leading to SEAP expression and secretion.
- **Raji-CD70 Ligand Cells:** Raji B-lymphoma cells that stably express the human CD70 ligand, which serves as the activator for the Jurkat-CD27 Reporter Cells.

Jurkat-CD27 Reporter Cells and Raji-CD70 Ligand Cells were mixed at a 1:1 ratio, total cell density 5×10^5^ cells/mL and resuspended in various ATPS compositions (prepared in RPMI-1640 medium supplemented with 10% FBS and 1% P/S), solutions containing FUS (2500 nM, 5000 nM, in RPMI-1640 medium), or control media (complete RPMI-1640). The positive control was co-culture in complete RPMI-1640 without ATPS/FUS.

Cells were incubated at 37°C in a humidified atmosphere containing 5% CO_2_. At indicated time points (4, 12, 24, 48, 72 h), SEAP activity in the cell culture supernatant was measured. Briefly, 20 *µ*L of supernatant was collected and added to 180 *µ*L of QUANTI-Blue^™^ Solution (InvivoGen, Cat #rep-qbs) in a 96-well plate. The plate was incubated at 37°C for 1 hour, and the absorbance at 655 nm was measured using a microplate reader (CytoFLEX, Beckman). The absorbance at 655 nm is directly proportional to the SEAP activity, reflecting NF-*κ*B activation and CD27/CD70 signaling.

### Illustrations and Conceptual Videos

All schematic illustrations and conceptual animations presented in the main figures, graphical abstract, and supplementary videos were meticulously designed and rendered using **Cinema 4D (Maxon)**. This professional 3D computer graphics software was utilised to create high-quality, scientifically accurate, and visually compelling representations of the proposed molecular mechanisms, experimental setups, and dynamic cellular interactions.

### Quantification and Statistical Analysis

Statistical analyses were performed using GraphPad Prism 10. For comparisons between two groups, a two-tailed unpaired Student’s t-test was used. For multiple comparisons, one-way ANOVA followed by Tukey’s post-hoc test was used. P values *<* 0.05 were considered statistically significant (*P <* 0.05, \*\**P <* 0.01, \*\*\**P <* 0.001, \*\*\*\**P <* 0.0001).

### Coarse-Grained Particle-Field Hybrid Model

We develop a coarse-grained particle–field hybrid model to simulate cell aggregation dynamics in phase-separating fluids, incorporating many-body hydrodynamic interactions (HI) through the fluid particle dynamics (FPD) method [6, 47, 48, 60]. The FPD method enables efficient evaluation of both far-field and near-field HI [61–64].

Within this framework, each cell particle is modeled as a viscous fluid particle represented by a smooth interfacial profile,

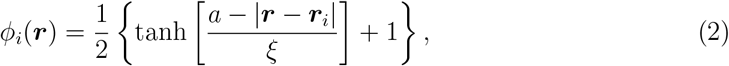

where *a* is the particle radius, *ξ* the interfacial thickness, and ***r***_*i*_ the position vector of the th particle in a cubic periodic box of size *L*. The viscosity field is then

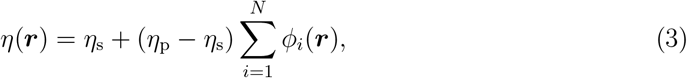

where *η*_s_ and *η*_p_ are the viscosities of the solvent and particles, respectively, and *N* is the total number of particles..

We employ the following free energy functional for the binary fluid mixture containing particles [60]:

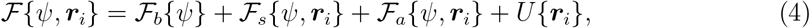

where the order parameter *ψ*(***r***) is the composition field.

#### Bulk free energy

The first term is the bulk Ginzburg–Landau (GL) free energy functional,

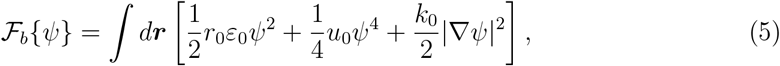

where *ε*_0_ = (*T* − *T*_c_)*/T*_c_ is the reduced temperature. For *r*_0_*ε*_0_ *<* 0 and *u*_0_ *>* 0, the system phase-separates into phases *A* and *B* with equilibrium compositions

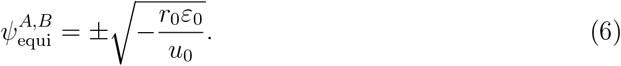

The quartic term stabilises the order parameter against unbounded growth.

#### Surface wetting energy

Particle–fluid wetting is described by

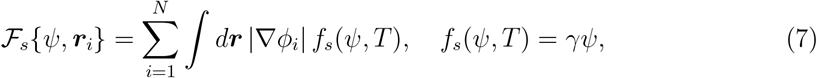

where *γ* is the coupling constant; *γ <* 0 corresponds to preferential wetting of phase *A* [47].

#### Particle impermeability

To enforce impermeability of particles to the order parameter, we introduce

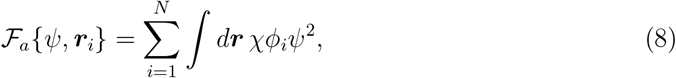

with *χ >* 0 ensuring *ψ* ≈ 0 inside the particles.

#### Direct particle interactions

Excluded volume interactions are modeled with the Weeks– Chandler–Andersen (WCA) potential,

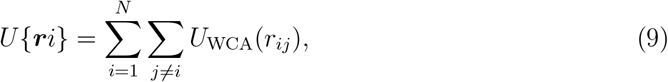

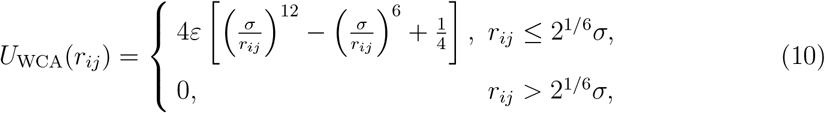

where *r*_*ij*_ = |***r***_*i*_ − ***r***_*j*_|, *σ* is the effective diameter, and *ε* the repulsion strength.

#### Hydrodynamics

The flow field ***v*** obeys the incompressible Navier–Stokes equation,

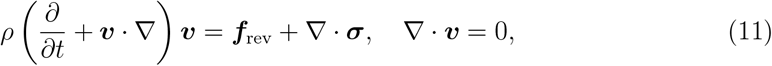

with density *ρ*, viscous stress tensor

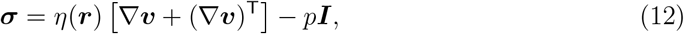

and pressure *p* enforcing incompressibility (**∇** · ***v*** = 0). The reversible force density is

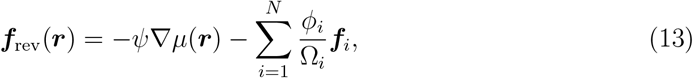

ensuring energy dissipation *d*ℱ/*dt* ≤ 0 while conserving momentum [60]. Here *µ* = δℱ*/*δ*ψ* is the chemical potential, ***f***_*i*_ = ∂ℱ*/*∂***r***_*i*_ is the force on particle *i*, and Ω_*i*_ = *d****r*** *ϕ*_*i*_ its effective volume.

#### Composition dynamics

The order parameter evolves as

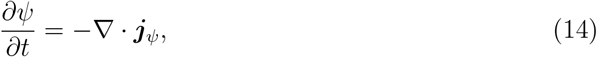

with flux

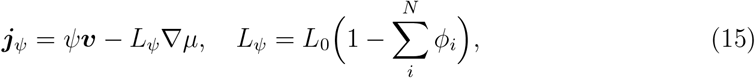

where *L*_0_ is a positive constant, and *L*_*ψ*_ vanishes inside particles.

#### Numerics

We solve Eq. (11) using the Marker-and-Cell (MAC) method[65] on a staggered grid with periodic boundaries. Particle positions are updated by

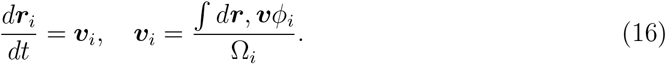

#### Quasi-2D confinement with planar walls

For modeling cells under quasi-2D confinement, we introduce planar walls located at *z* = 0 and *z* = *h*, described by

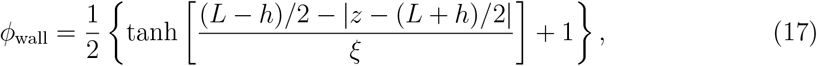

where the walls, having a high viscosity *η*_p_, effectively behave like a solid.

#### Wall–particle interactions

Three additional terms are incorporated into the free energy ℱ{*ψ*, ***r***_*i*_}. First, the WCA potential 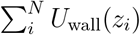 between particles and walls is added to the potential energy *U* {***r***_*i*_} to prevent particle penetration. The potential is defined as

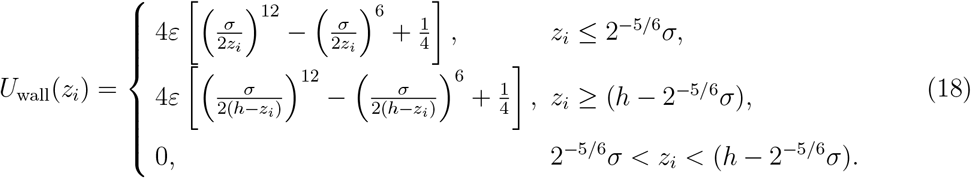

#### Wall impermeability

Second, the term

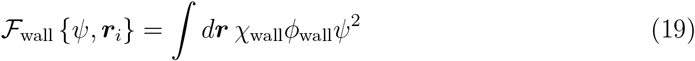

is added to ℱ_*a*_ {*ψ*, ***r***_*i*_} to ensure *ψ* ≈ 0 inside the walls, analogous to the treatment for particles [Eq. 8].

#### Wall wetting

Third, the affinity of the minority phase for the walls is described by the wetting term

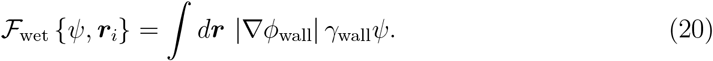

We set *γ*_wall_ = 0 for neutral walls and *γ*_wall_ = −4 for preferential wetting walls.

#### No-slip condition and external flow

To impose a no-slip boundary condition, we enforce ***v*** = 0 on the walls by adding a frictional body force

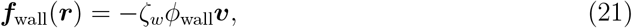

which suppresses flow penetration into the walls [66, 67]. Additionally, a constant drag force

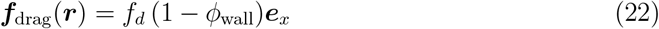

is applied along the *x*-direction to generate a pressure-driven Poiseuille flow with a parabolic velocity profile in the absence of particles.

#### Simulation parameters

We set the length unit as the lattice spacing *l*_0_, the time unit as 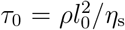, *a* = 3.2*l*_0_, *ξ* = *l*_0_, particle diameter *σ* = 2*a* + *ξ* = 7.4*l*_0_, *η*_p_ = 50*η*_s_, *ε* = 100, *L*_0_ = 1, *L* = 256*l*_0_ = 34.59*σ*, and time step Δ*t* = 2.5 × 10^−3^*τ*_0_. For no-slip walls, we use ζ_*w*_ = 50 and wall separation *h* = 150*l*_0_. The drag force *f*_*d*_ is varied as *f*_*d*_ = 0, *f*_*d*_ = 5 × 10^−6^, and *f*_*d*_ = 5 × 10^−5^ to achieve different flow rates. We set *χ* = 2 and *χ*_wall_ = 4 to ensure particle/wall impermeability. The surface coupling strength is varied from *γ* = −8 (strong phase affinity) to *γ* = −4 (moderate affinity) and *γ* = −1 (weak affinity) to examine the role of particle–fluid wetting. By tuning 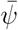 from 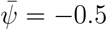 to 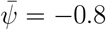 (moderately asymmetric) and 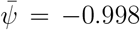 (highly asymmetric), we vary the composition ratio 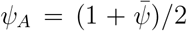 of phase *A* from *ψ*_*A*_ ~ 25% to *ψ*_*A*_ ~ 0.1%, thereby controlling the minority phase fraction.

#### Free energy parameters

For the GL free energy, we set *r*_0_*ε*_0_ = −0.85, *u*_0_ = 1, and *k*_0_ = 1 in 3D and quasi-2D simulations. This choice allows us to probe the one-phase regime 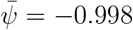, reproducing experimental observations where cell aggregation arises even without bulk phase separation. In 2D simulations, we set *r*_0_*ε*_0_ = −1, *u*_0_ = 1, and *k*_0_ = 1.

#### Initial conditions

All simulations begin from a homogeneous initial state where particles are randomly distributed in space. Thermal noise is neglected, which is a reasonable approximation for biological cells of size *σ* ≈ 10 *µ*m.

### Number of Particles in a Cluster *N*_*c*_

We employed a cluster analysis algorithm implemented in Ovito [68] to compute the number of particles *N*_*c*_ in a cluster. The algorithm assigns particles to the same cluster when their separation distance is less than a cutoff distance *r*_cut_. To capture connectivity through the minority phase, our analysis includes both the grid points with *ψ*(***r***) *>* 0 and the grid points inside each particle as pseudo-particles. This ensures that only phase-connected particles are classified as belonging to the same cluster (we set *r*_cut_ = *l*_0_). Thus, we obtain the average cluster size *N*_*c*_ connected through the minority phase.

### Average Distance to Walls *d*_wall_

To quantify wall-directed particle aggregation in quasi-2D simulations, we computed the minimum perpendicular distance (along the *z*-axis) between each particle and the confining walls (*z* = 0 and *z* = *h*) at each time *t*. The values were averaged over all particles to obtain the mean particle–wall distance *d*_wall_.

### Cluster Participation Ratios *P*_1_ and *P*_2_ for Binary Systems

To study mixed populations of two cell types with distinct wetting affinities, we assigned surface coupling strengths *γ*_1_ (particle number *N*_1_ = *N/*2) and *γ*_2_ (particle number *N*_2_ = *N/*2). We then counted the number of *γ*_1_-type particles (*n*_1_) that clustered via the minority phase and defined their participation ratio as *P*_1_ = *n*_1_*/N*_1_. Analogously, the participation ratio *P*_2_ = *n*_2_*/N*_2_ was defined for *γ*_2_-type particles (*n*_2_ being the number incorporated into *γ*_1_-type clusters). Thus, *P*_1_ and *P*_2_ quantify the aggregation propensities of the two cell types.

## Data availability

All original code has been deposited at Mendely under the https://data.mendeley.com/drafts/wgb3d62gm8. The data that support the findings of this work are available within the article and the supplementary information. The raw data will be shared by the corresponding authors upon reasonable request.

## Code Availability

The code and analysis scripts used in this study are available from the corresponding authors upon reasonable request.

## Acknowledgements

We acknowledge the support on imaging from The Fifth Affiliated Hospital of Sun Yat sen University (Molecular Imaging Center) and Guangdong Institute of Intelligence Science and Technology, Hengqin (Platform for Structure and Functional Technology of Neural Network). We also thank Professor Shizhong Luo and his team for their generous help in providing Recombinant CCL5 protein. C.M. expresses gratitude to Mrs. Huihui Liu, Dr. Daping Xie, Dr. Zhencheng Liao, and Dr. Yiming Niu for their invaluable technical assistance. This study was financially supported by the Science and Technology Development Fund, Macao SAR (FDCT, No. 0001/2021/AKP, 0024/2023/AFJ, 0209/2024/AGJ, 0031/2023/ITP1, and 005/2023/SKL), the National Natural Science Foundation of China (NSFC, No. 32361163656, 32022088, and 32230056), the Natural Science Foundation of Jiangsu Province (BM2023008), the Guangdong Provincial Committee for Basic and Applied Basic Research (EF2023-00209-ICMS), and the University of Macau (MYRG-GRG2023-00136-ICMS-UMDF, MYRG-GRG2024-00189-ICMS-UMDF, and MYRG-CRG2023-00009-IAPME), as well as the Zhuhai UM Science & Technology Research Institute (CP-102-2024). J.Y. is supported by the startup fund provided by HKUST(GZ) and the National Natural Science Foundation of China (Grant No. 22503076). J.Y. thanks the HPC AI Intelligent Computing Platform at HKUST(GZ) for providing the GPU resources for simulations. H.T. acknowledges support from the Grant-in-Aid for Specially Promoted Research (JSPS KAKENHI Grant No. JP20H05619) from the Japan Society for the Promotion of Science (JSPS).

## Author contributions

Conceptualisation: A.W., H.T., J.Y., and C.W.; Methodology: A.W., D.Y., and J.Y.; Investigation: A.W., D.Y., H.Z., and J.Y.; Writing – Original Draft: A.W., H.T., J.Y., and C.W.; Writing – Review & Editing: A.W., H.T., J.Y., V.P., L.D., and C.W.; Funding Acquisition: H.T., J.Y., and C.W.; Resources: J.Y. and C.W.; Supervision: C.W., J.Y., H.T., and V.P.

## Additional Information

Requests for further information and resources should be directed to and will be fulfilled by the lead contact, Dr. Chunming Wang.

**Extended Data Fig. 1.**
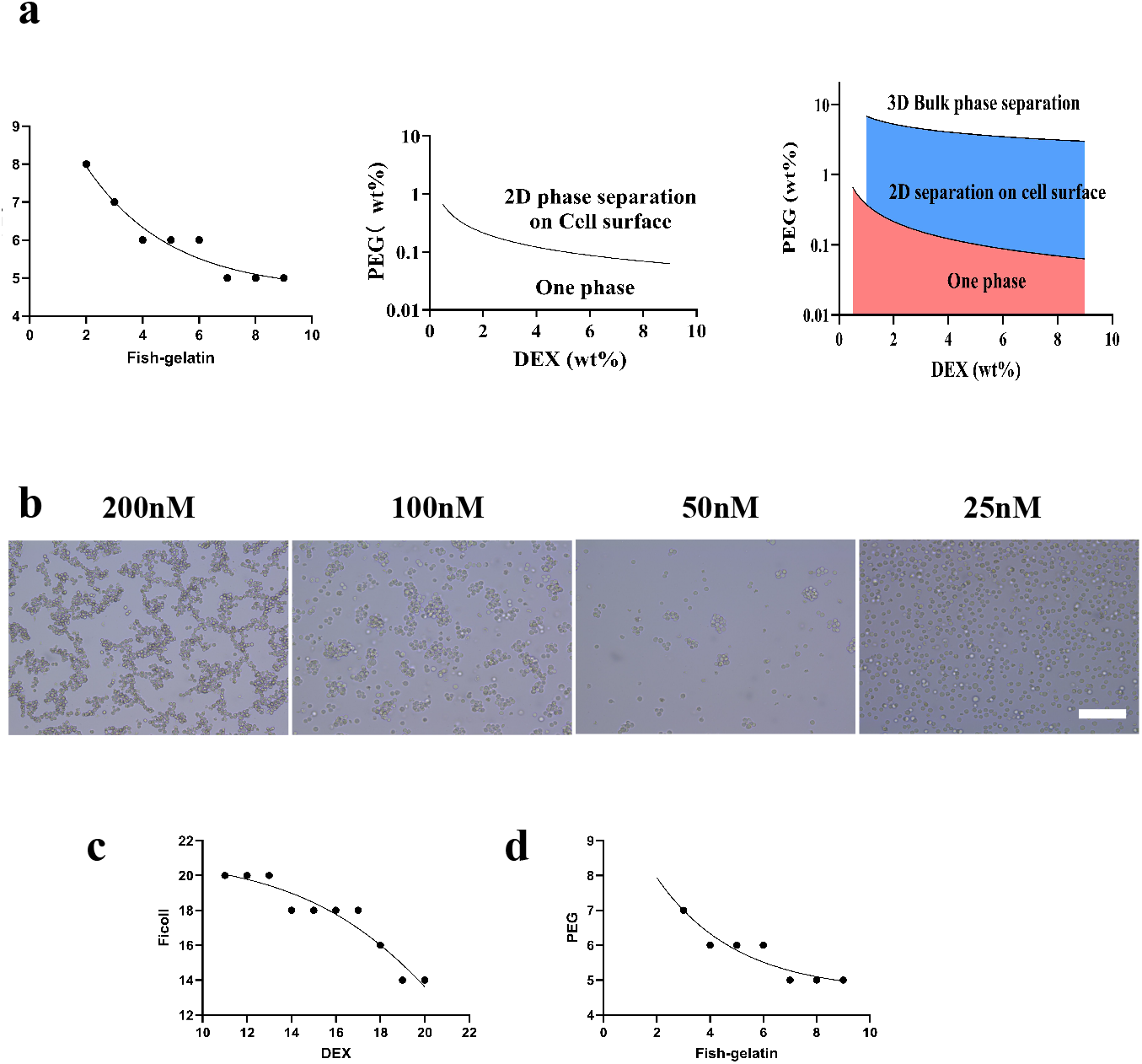
Experimental phase diagram and concentration dependence of aggregation. **a** Experimental binodal and bulk-one phase curve for the DEX/PEG ATPS and a combined diagram illustrating the regimes of surface-induced 2D separation below the bulk threshold. **b** Brightfield images showing that cell aggregation is dependent on Dextran concentration in bulk one-phase concentration. Scale bar, 100 *µ*m. **c** Phase diagram of the Dextran-Ficoll ATPS system. The binodal curve delineates the coexistence region where two distinct aqueous phases separate, representing the critical concentrations of Dextran and Ficoll for liquid-liquid phase separation. **d** Phase diagram of the PEG-Fish Gelatin ATPS system.

**Extended Data Fig. 2.**
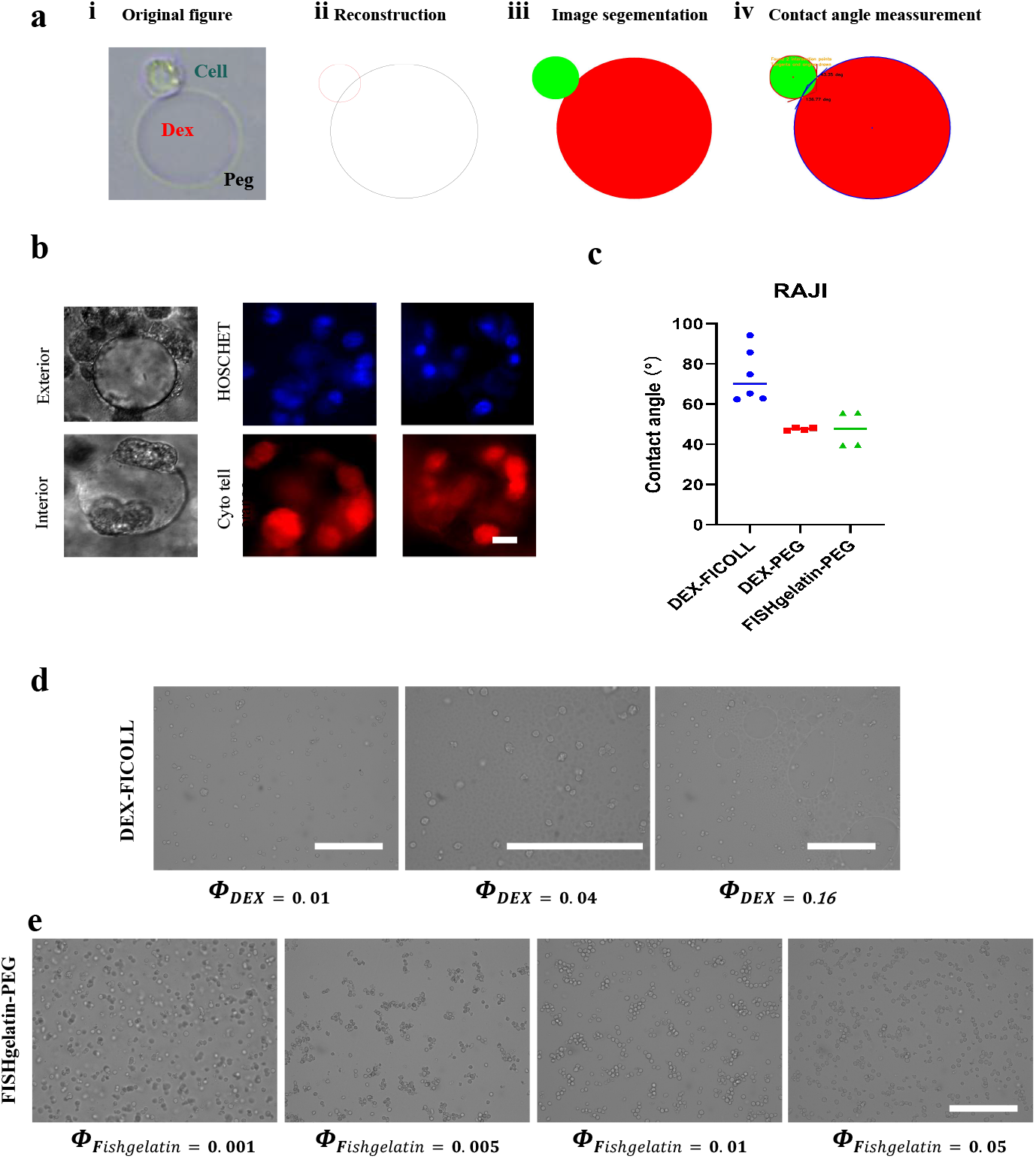
Methodology and controls for wettability analysis. **a** Image analysis pipeline for contact angle, *θ*, measurement. **b** Fluorescence microscopy images showing the localisation of Jurkat cells at the DEX/PEG interface. Scale bar, 10 *µ*m. **c** Contact angle data for Raji cells. **d** Negative control: negligible aggregation is observed in the non-wetting DEX-FICOLL system across various compositions (*ϕ*_DEX_). Scale bar, 100 *µ*m. **e** Generality control: robust cell aggregation is observed in the wetting FISHgelatin-PEG system across various compositions (*ϕ*_DEX_). Scale bar, 100 *µ*m.

**Extended Data Fig. 3.**
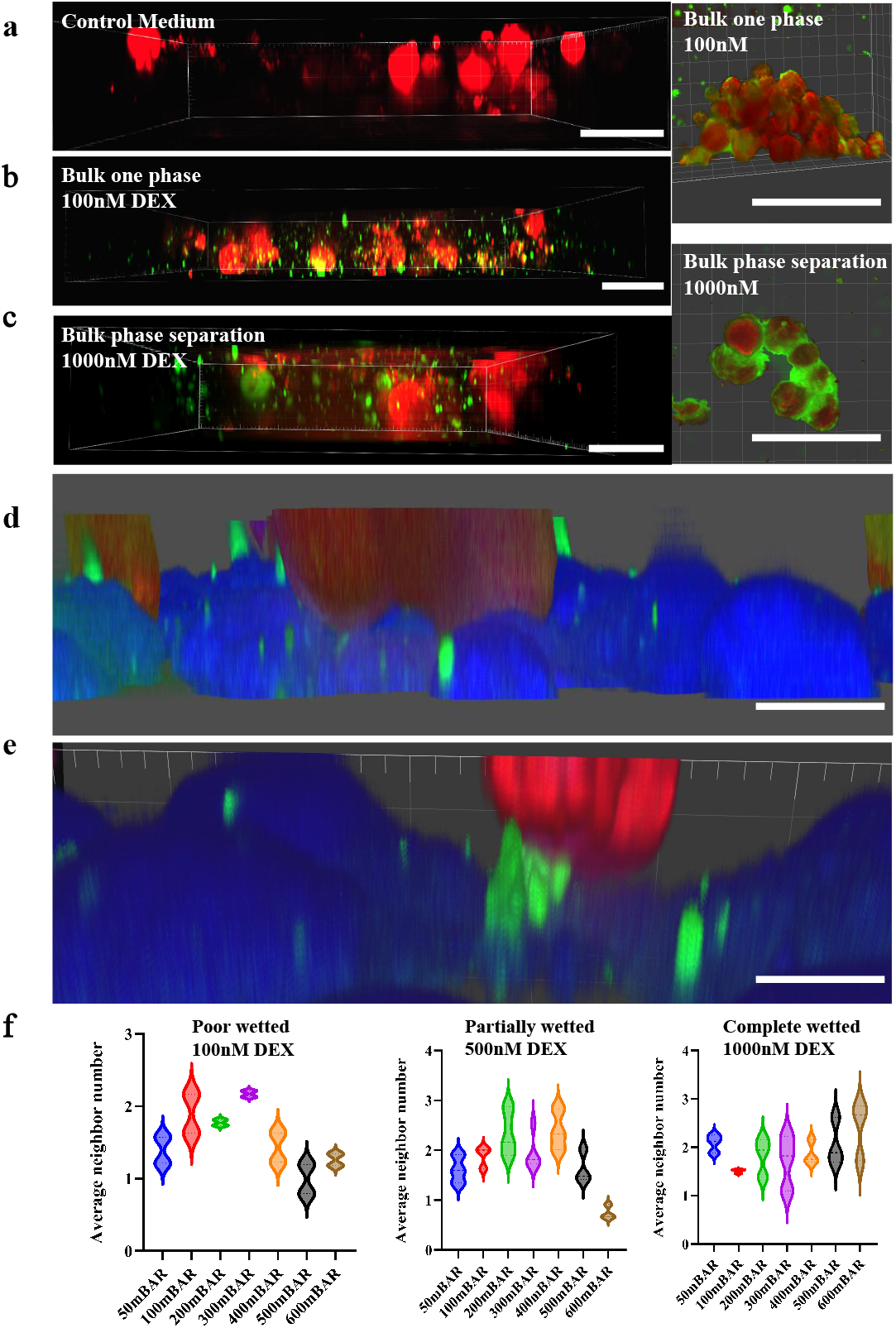
Effect of shear flow on cell aggregation under different wetting conditions. **a–c** 3D confocal reconstructions of Jurkat-Petri dish aggregates under ‘Non-wetted (Control medium)’, ‘poorly wetted (100nM DEX)’, and ‘completely wetted (1000nM dex)’ conditions, respectively. Scale bar, 50 *µ*m. **d–e** Side-view projections of Jurkat-HCMEC aggregates cell. Scale bar, 10 *µ*m. **f** Quantitative analysis reveals a non-monotonic relationship between shear pressure and aggregation. This effect, where an optimal pressure enhances aggregation, is prominent in the ‘Poor wetted’ (100 nM DEX) and ‘Partially wetted’ (500 nM DEX) conditions but not in the ‘Completely wetted’ (1000 nM DEX) state.

**Extended Data Fig. 4.**
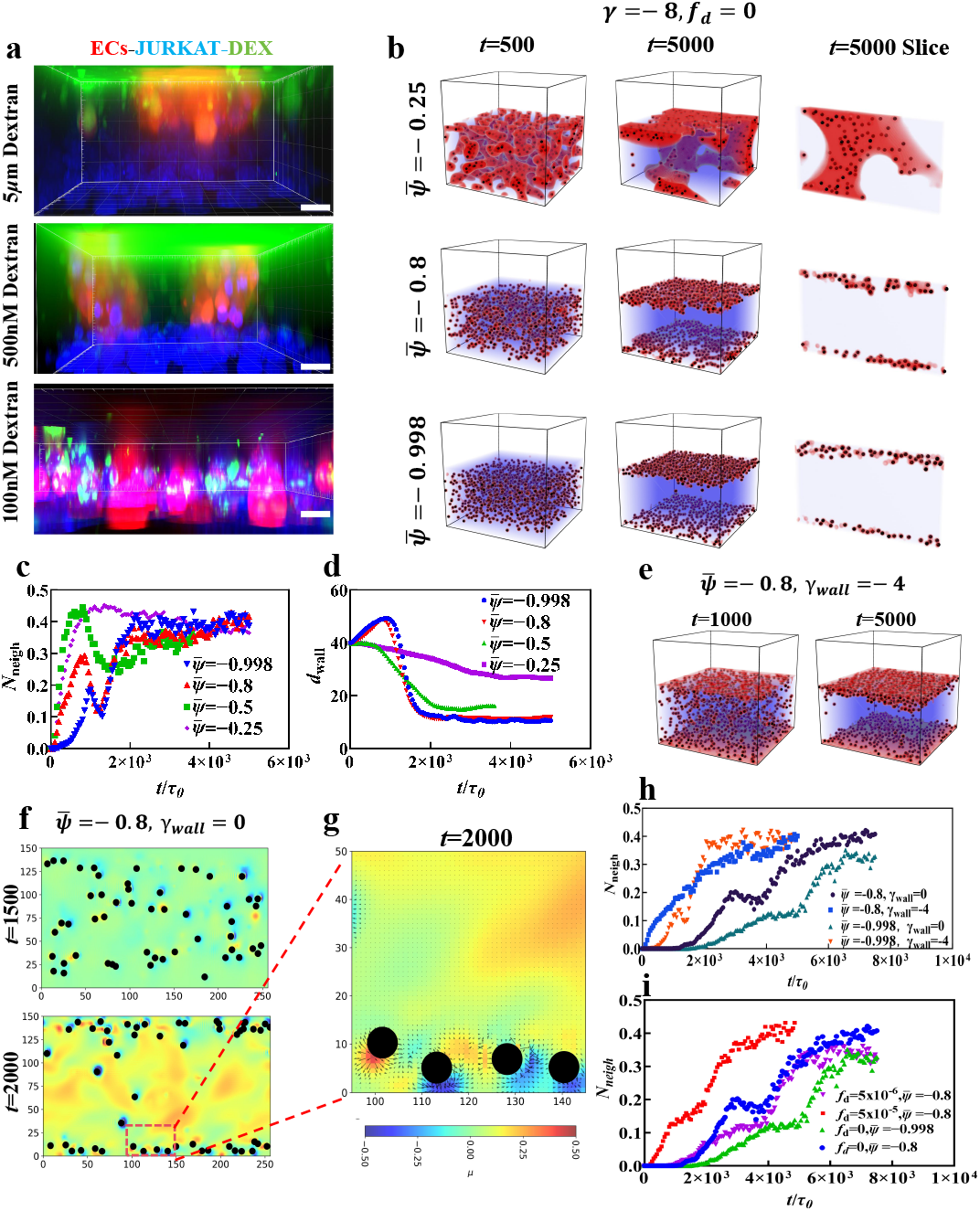
Adhesion of suspended cells to surfaces. **a** Experimental co-culture of Jurkat (red) and endothelial cells (green) with DEX (blue) at high (5 *µ*M), intermediate (500 nM), and low (100 nM) concentrations. Scale bars, 10 *µ*m. **b** 3D simulations of particle aggregation (*ϕ*_Cell_ = 2.5%) on neutral walls for different average compositions 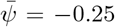, −0.8, and −0.998 under *f*_*d*_ = 0 (no external flow). **c** Temporal evolution of the average neighbor number *N*_neigh_ for the simulations shown in **b. d** Temporal evolution of the average particle-to-wall distance *d*_wall_ for the simulations shown in **b. e** 3D simulation snapshots of particle adhesion to an attractive wall with wetting affinity *γ*_wall_ = −4 at 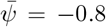. **f** Spatiotemporal evolution of the chemical potential field for particles adhered to a neutral wall (*γ*_wall_ = 0) at 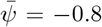. **g** Magnified view of **f**, showing the chemical potential *µ* and its negative gradient −∇*µ* (arrows). **h** Kinetics of *N*_neigh_ comparing different wall affinities (*γ*_wall_ = 0 vs. −4) and average compositions (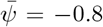 vs. −0.998). **i** Kinetics of *N*_neigh_ at 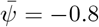 and 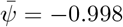, comparing the static case (*f*_*d*_ = 0) with different strengths of external flow (*f*_*d*_ *>* 0).

**Extended Data Fig. 5.**
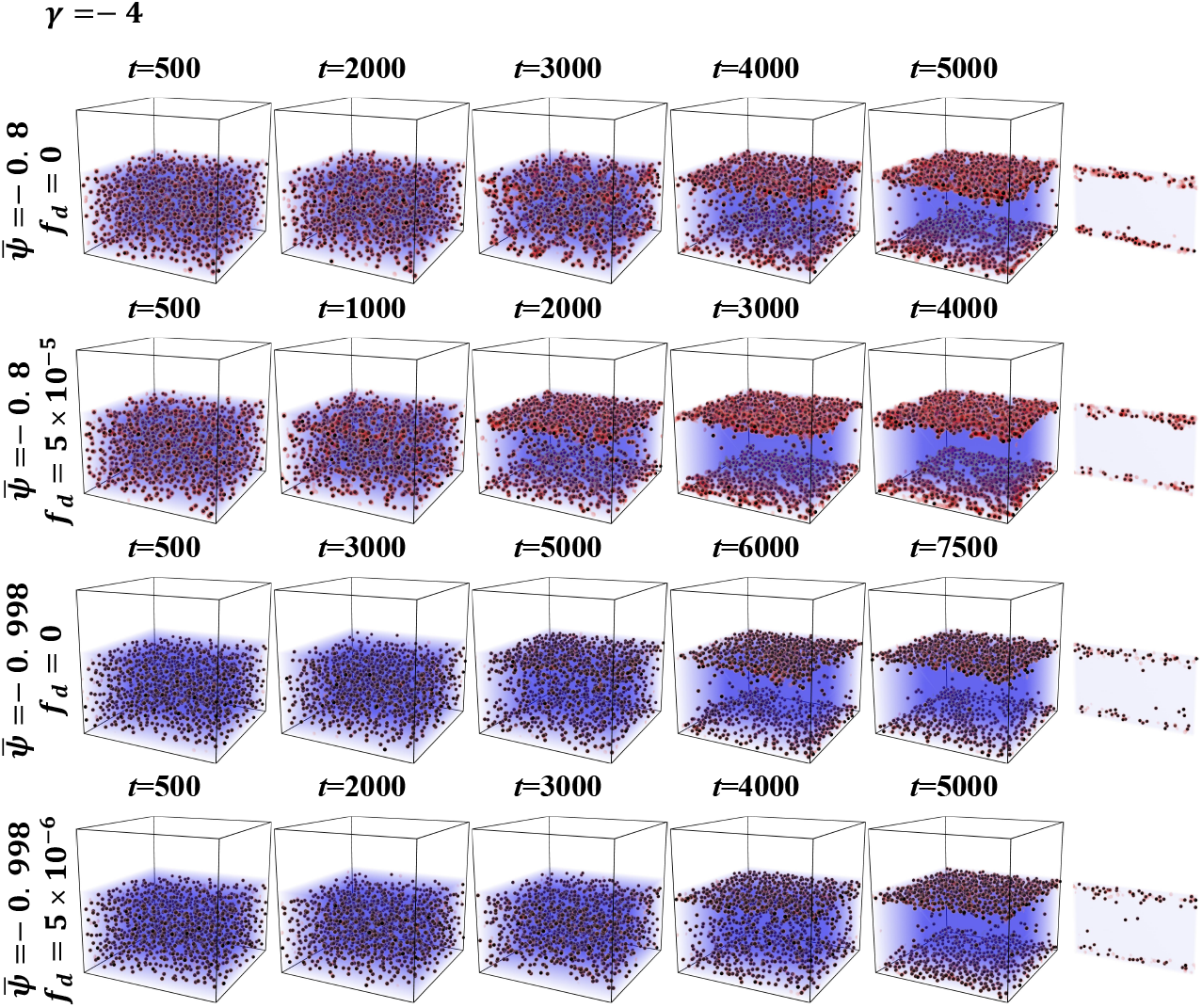
Quasi-2D simulations of cell adhesion to wall surfaces under intermediate wetting affinity (*γ* = −4). Adhesion to the wall surface occurs more slowly when the wetting affinity is weak compared to the case of strong wetting affinity (*γ* = −8).

**Extended Data Fig. 6.**
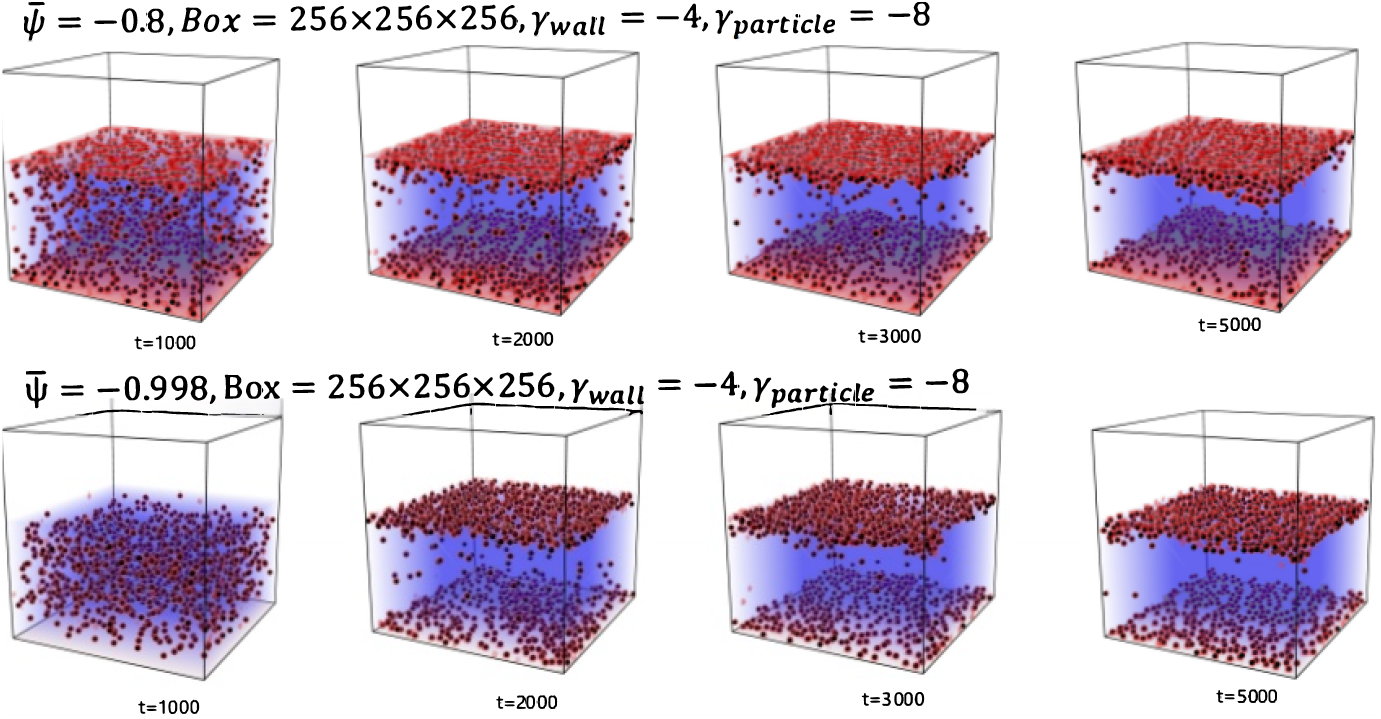
Quasi-2D simulations of cell adhesion to attractive wall surfaces (*γ*_wall_ = −4) under strong wetting affinity (*γ* = −8). A wall with an affinity for the minority phase results in faster adhesion compared to a neutral wall (*γ*_wall_ = 0; **Extended Data Fig**. 4). The wetting affinity (*γ*_wall_) also causes a minority-phase film to coat the wall, especially for the case of 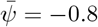.

**Extended Data Fig. 7.**
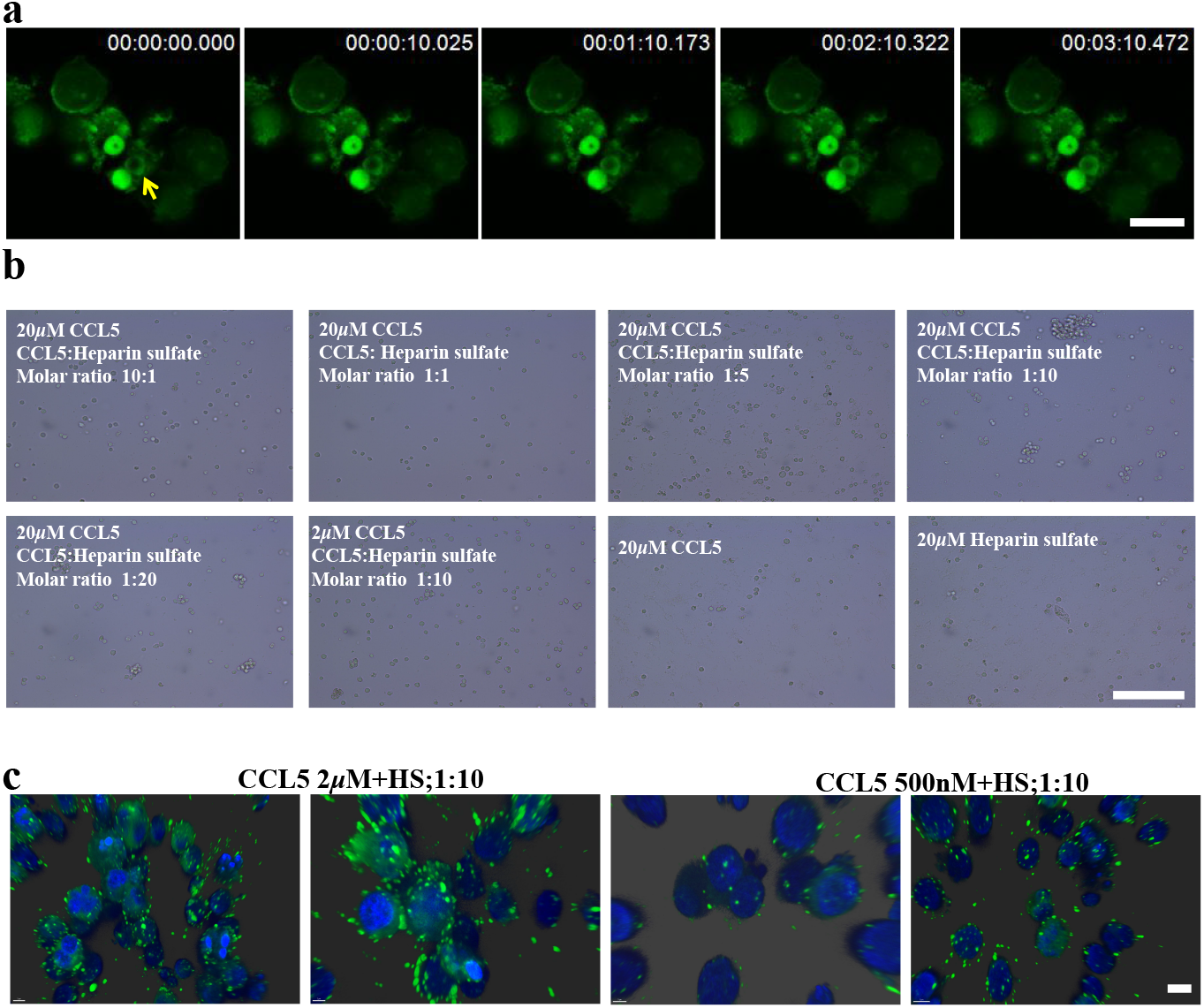
Cell aggregation induced by GAL3, CCL5/heparan sulfate. **a** Time-lapse showing liquid-like coalescence of 100nM GAL3 condensates on cell surface. **b** Bright-field images showing that aggregation is dependent on the CCL5:HS molar ratio. **c** Confocal images visualizing the condensates of CCL5/HS on the cell surface at different concentration. Scale bar, 10 *µ*m.

