## Supplementary material for "Wetting-mediated extracellular phase separation drives long-range cell adhesion": SI

**a**

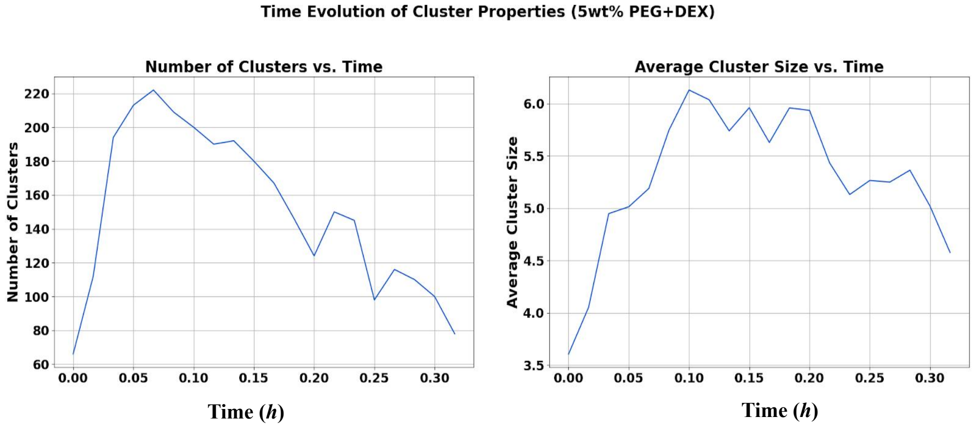


**b**

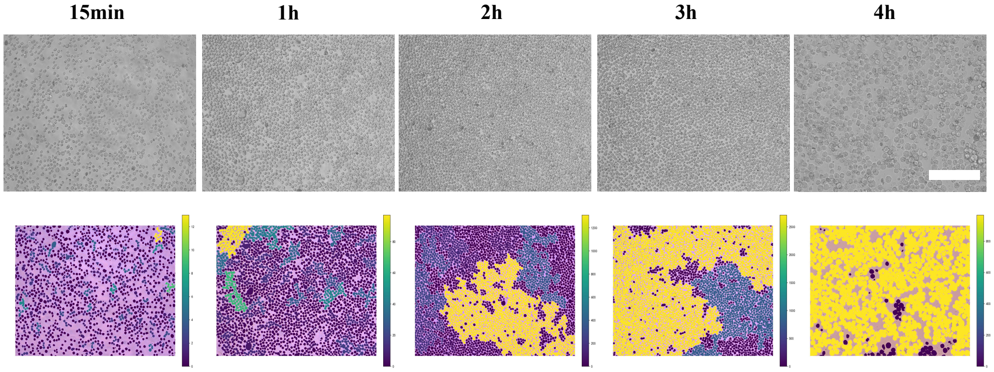


FIG. S1. **Kinetics of LLPS-driven vs. spontaneous cell aggregation. a** Quantification of Jurkat cell cluster properties over time in an ATPS with $\phi_{\mathrm{DEX}} = 0.01$. The number of clusters initially increases and then decreases, while the average cells per cluster follows the same trend. Scale bar, 100 $\mu$m. **b** Time-lapse microscopy of spontaneous Jurkat cell aggregation in a control single-phase system, demonstrating significantly slower kinetics compared to the LLPS-driven process. Purple indicates isolated cells, while yellow/green regions (as shown in the color map) represent increasingly connected or clustered cells. Scale bar, 100 $\mu$m.

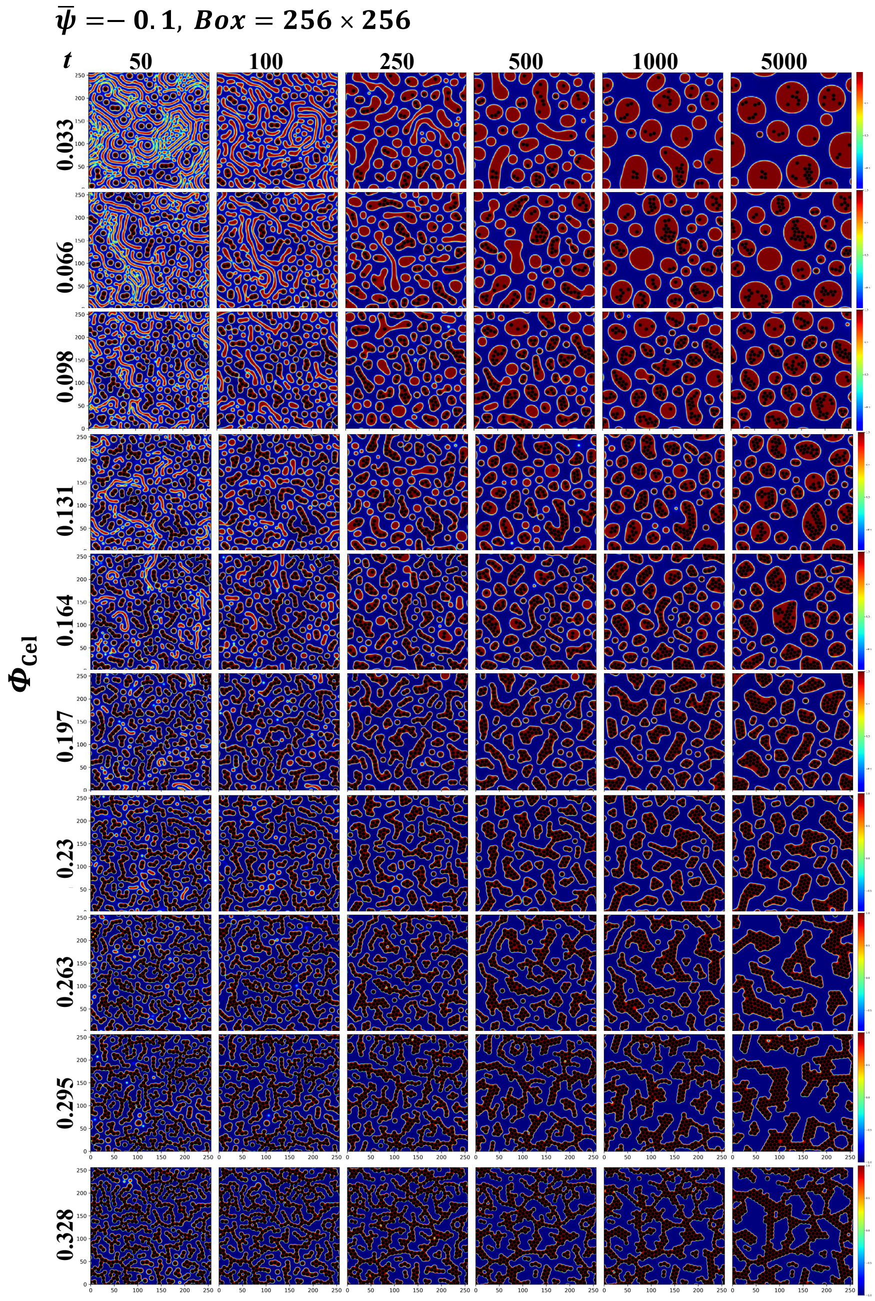


FIG. S2. **Simulated morphological evolution of particles in a phase-separating fluids with average composition $\bar{\psi} = -0.1$.** Time-series of 2D simulations showing the effect of cell occupation fraction, $\phi_{\text{Cell}}$, on morphology at $\bar{\psi} = -0.1$. The system exhibits round droplets (low $\phi_{\text{Cell}}$), branched clusters (intermediate $\phi_{\text{Cell}}$), and network-forming states (high $\phi_{\text{Cell}}$). The color map corresponds to the local $\psi$, with $\psi = 1$ and $\psi = -1$ representing the two distinct phases.

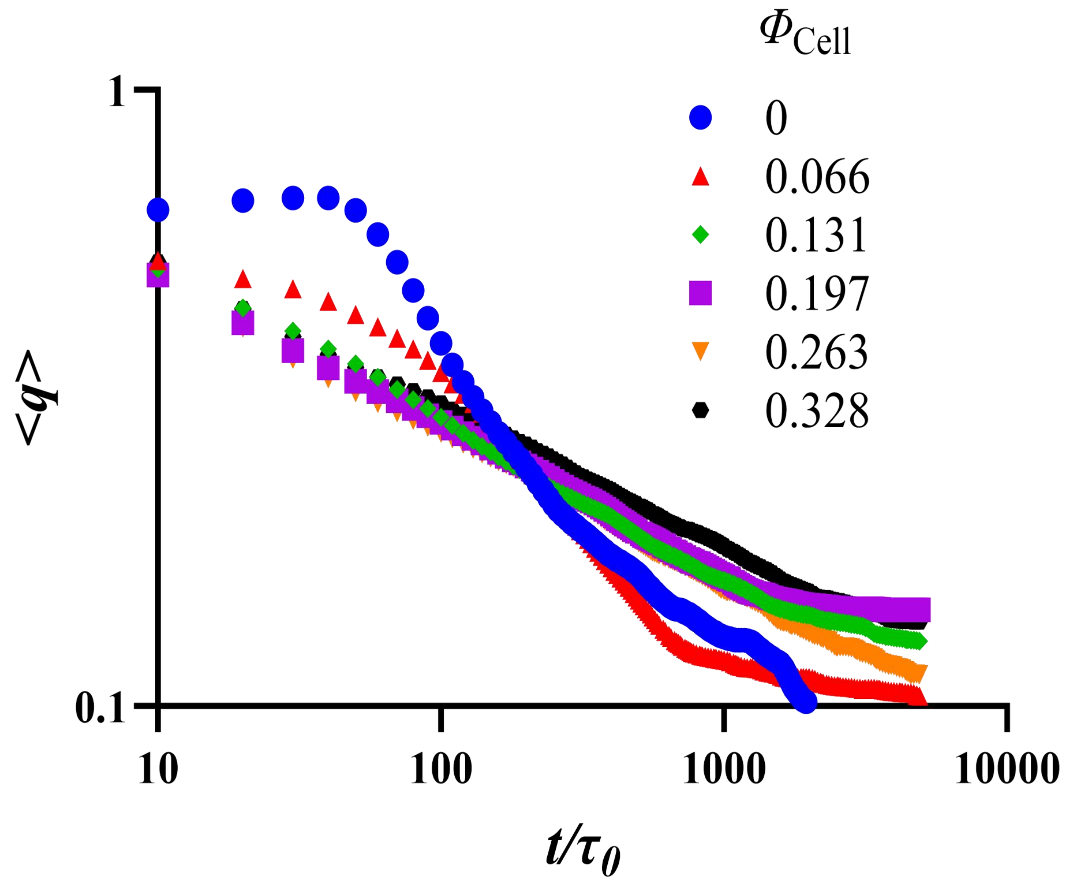


FIG. S3. **Quantitative analysis of coarsening dynamics at $\bar{\psi} = -0.1$.** Log-log plot of the average wave number, $\langle q \rangle$, versus time for the simulations shown in Fig. S2. The presence of particles (all non-blue curves) accelerates early-stage coarsening because of wetting effects but slows down the late stage compared to the particle-free system (blue curve), suggesting kinetic arrest.

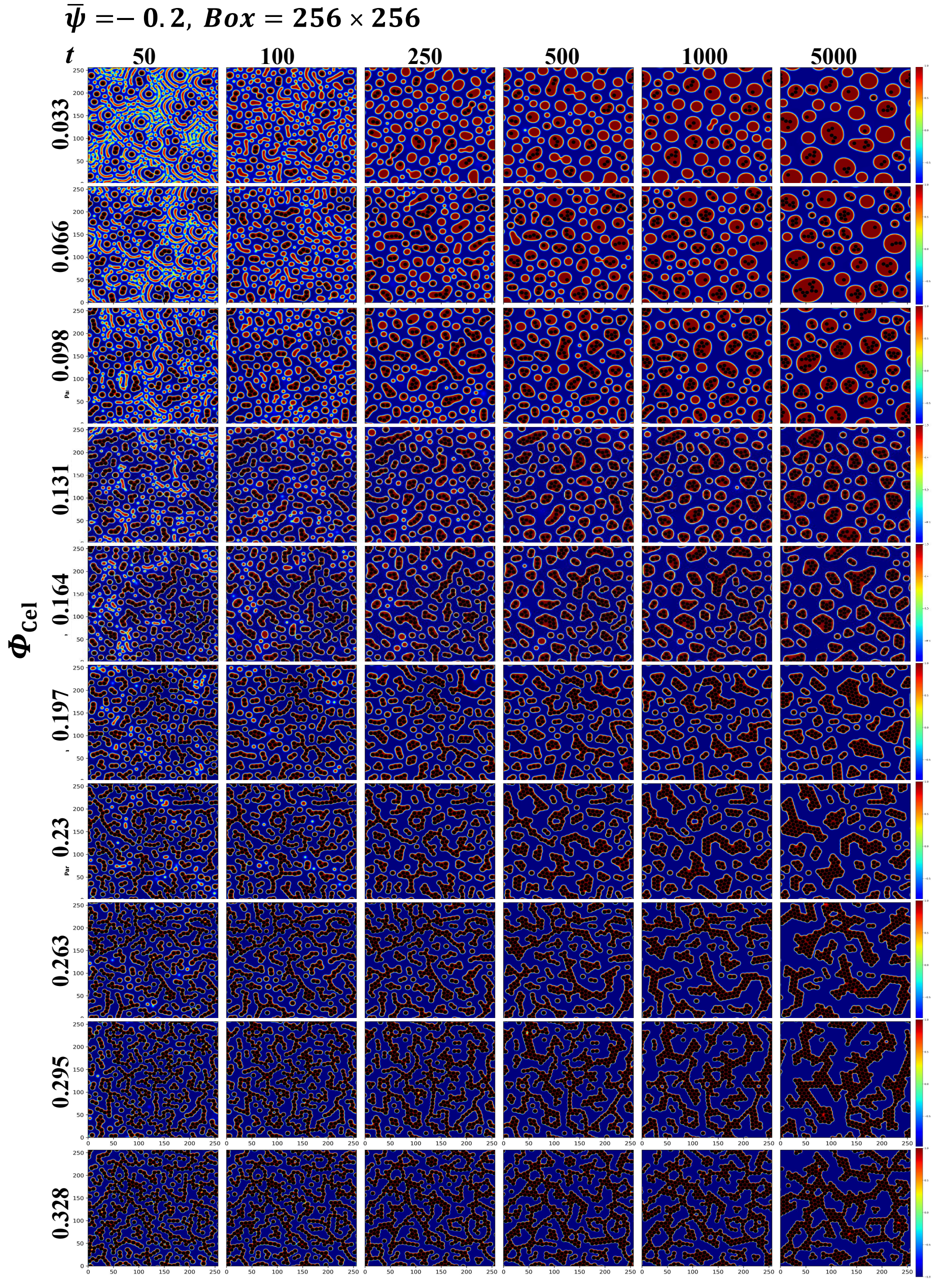


FIG. S4. **Simulated morphological evolution of particles in a phase-separating fluids with average composition $\bar{\psi} = -0.2$.**

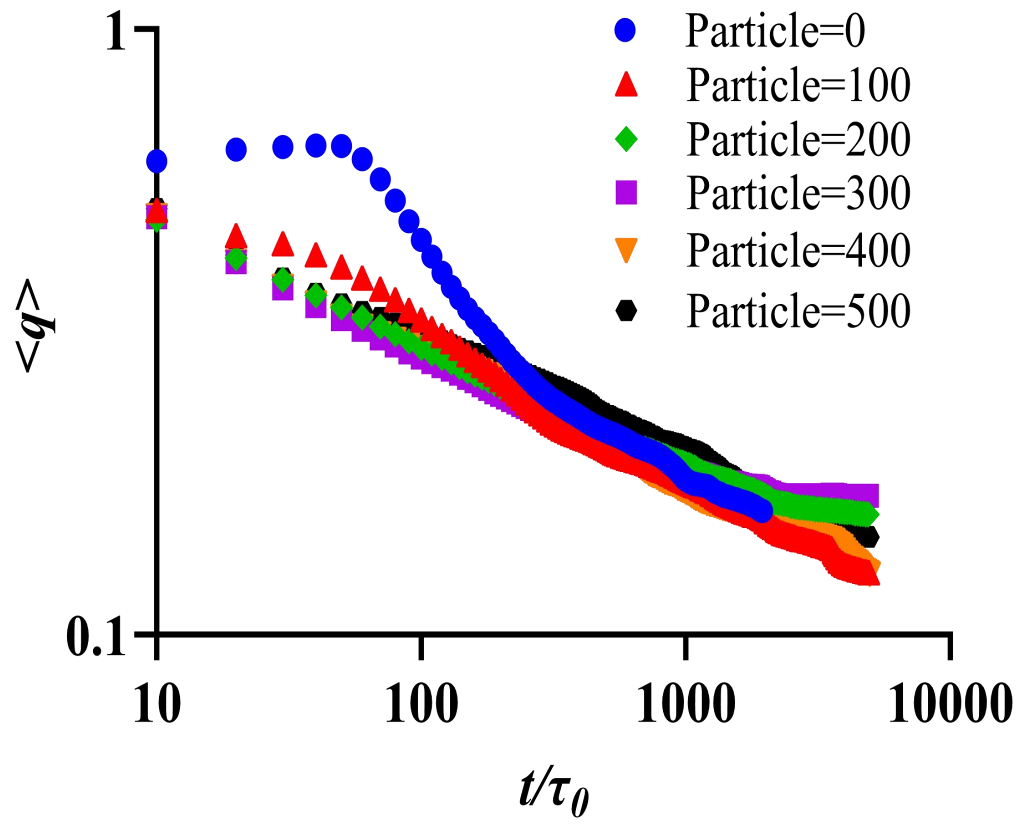


FIG. S5. **Quantitative analysis of coarsening dynamics at $\bar{\psi} = -0.2$.** Log-log plot of the average wave number, $\langle q \rangle$, versus time for the simulations in Fig. S4.

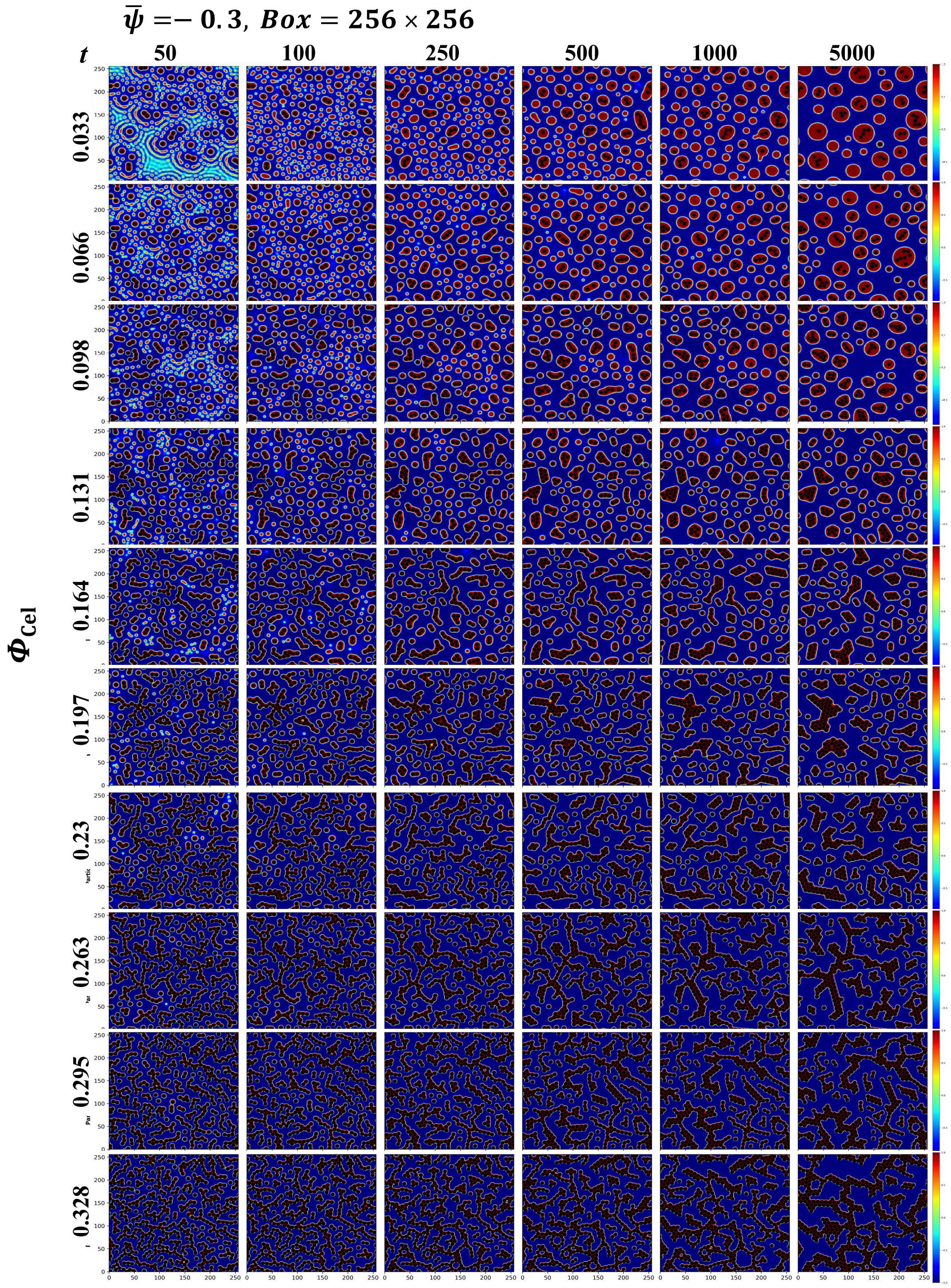


FIG. S6. **Simulated morphological evolution of particles in a phase-separating fluids with average composition $\bar{\psi} = -0.3$.**

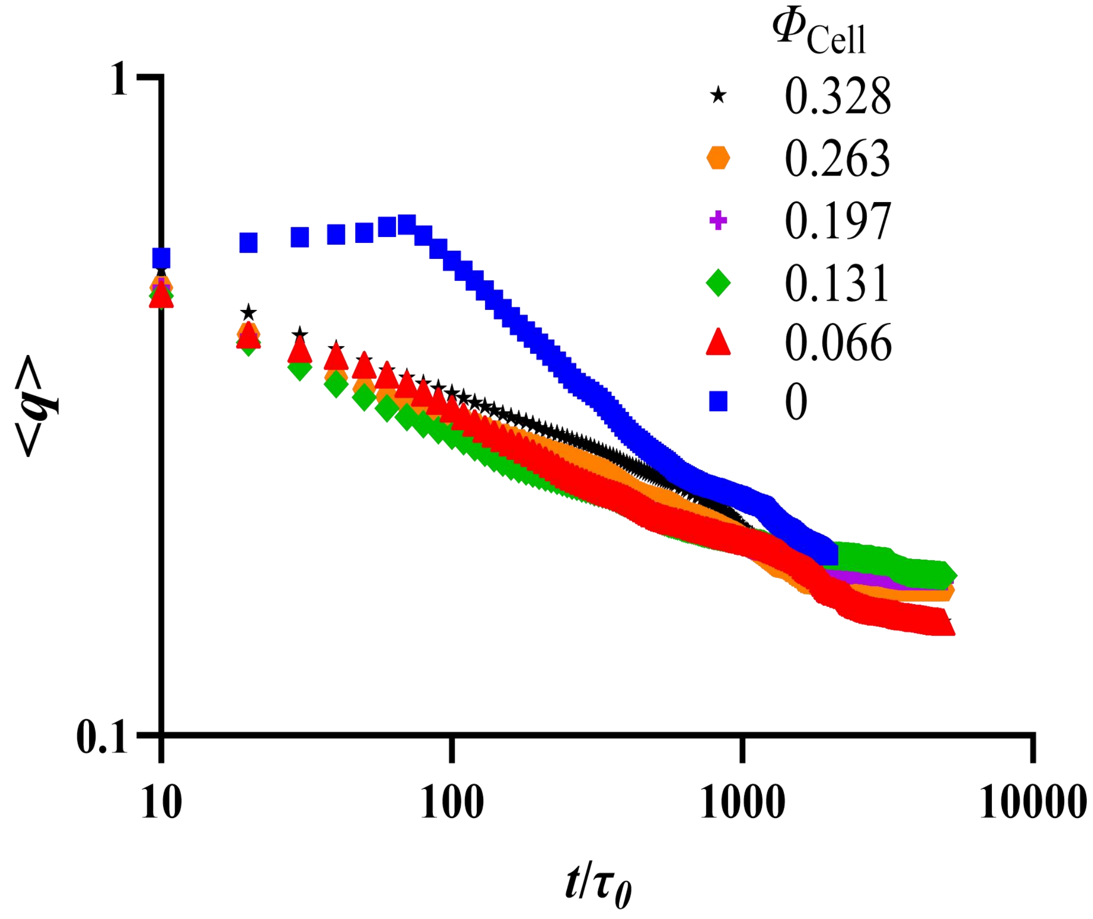


FIG. S7. **Quantitative analysis of coarsening dynamics at $\bar{\psi} = -0.3$.** Log-log plot of the average wave number, $\langle q \rangle$, versus time for the simulations in Fig. S6.

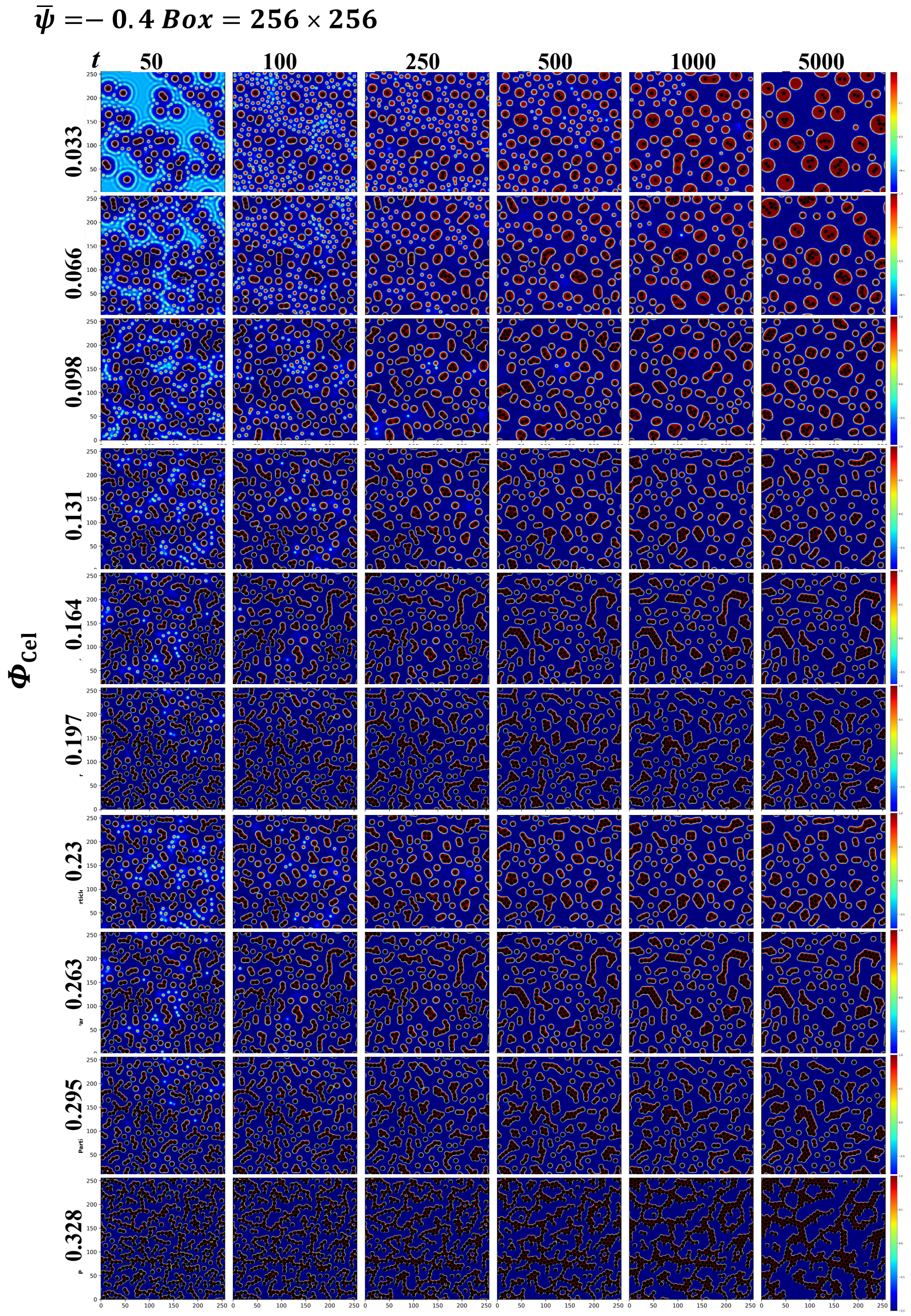


FIG. S8. **Simulated morphological evolution of particles in a phase-separating fluids with average composition $\bar{\psi} = -0.4$.** The trend qualitatively parallels the case of $\bar{\psi} = -0.1$. However, at a lower cell occupation fraction, the system begins to form elongated, branched clusters.

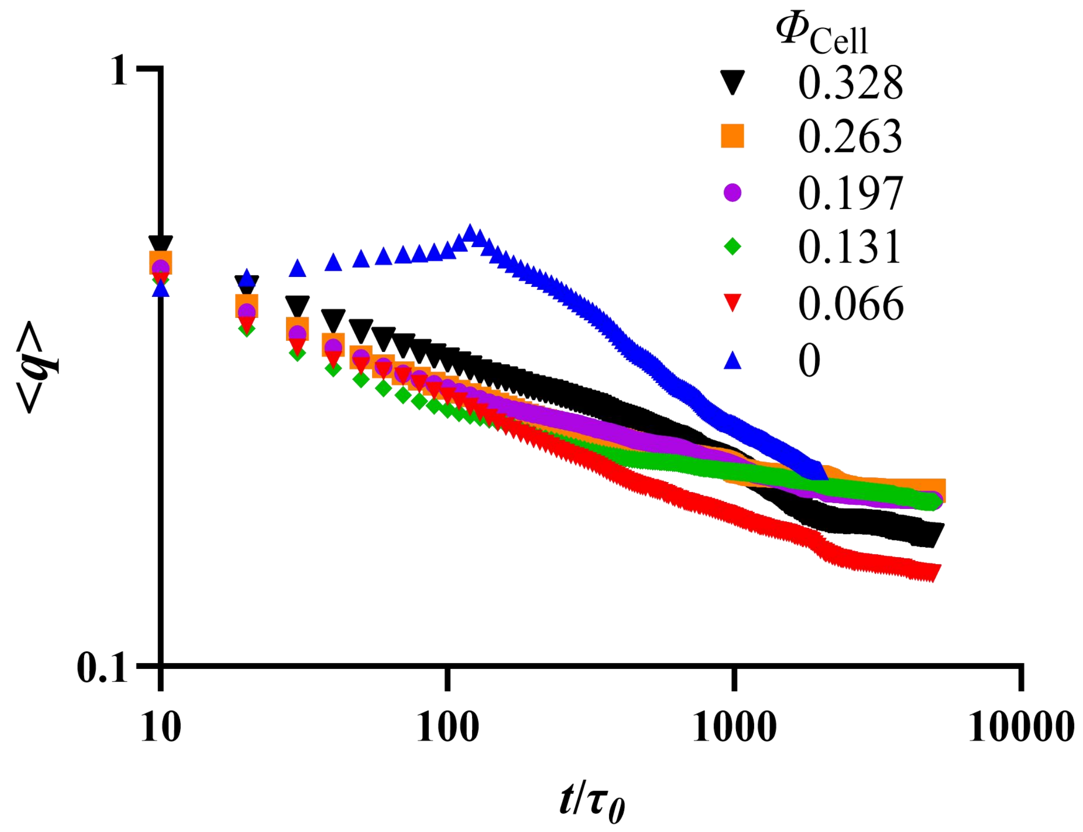


FIG. S9. **Quantitative analysis of coarsening dynamics at $\bar{\psi} = -0.4$.** Log-log plot of the average wave number, $\langle q \rangle$, versus time for the simulations in Fig. S8. The data confirms that particle aggregates kinetically arrest coarsening under $\bar{\psi} = -0.4$ as well.

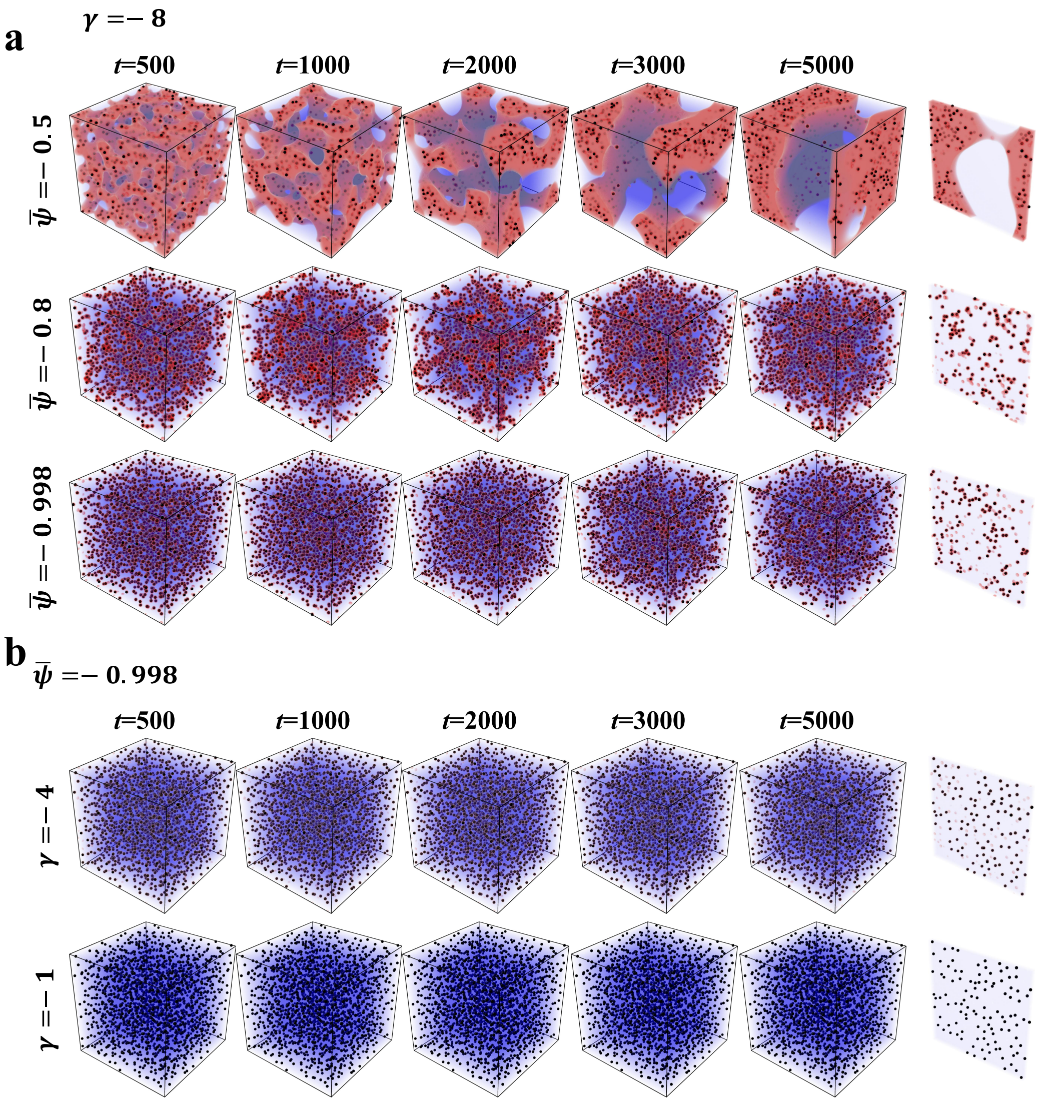


FIG. S10. **3D simulations on the effects of wetting phase volume and surface affinity. a** Effect of wetting phase volume ($\bar{\psi}$) at constant high affinity ($\gamma = -8$). **b** Effect of surface affinity ($\gamma$) at constant low phase volume ($\bar{\psi} = -0.998$), showing that high affinity is a prerequisite for aggregation.

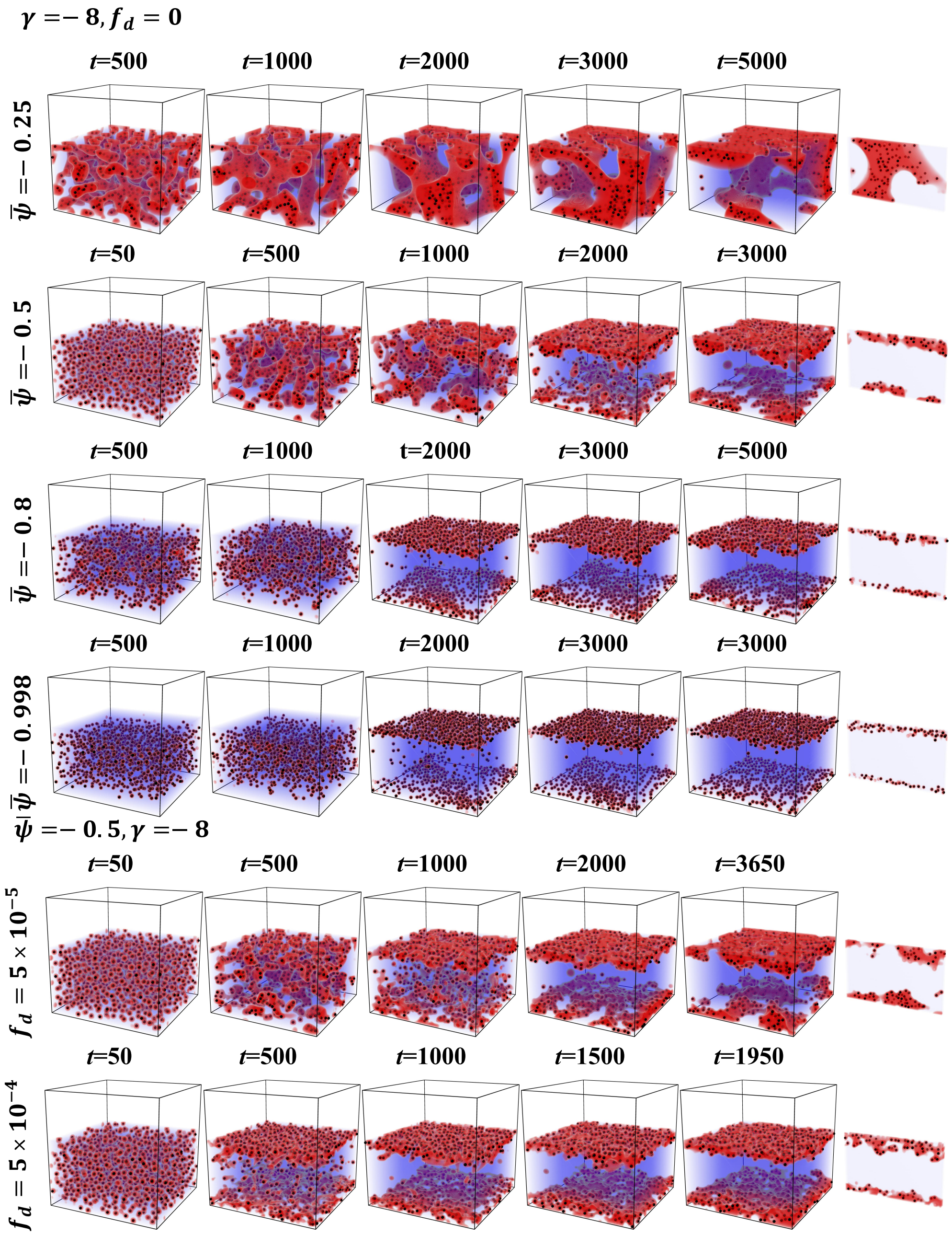


FIG. S11. **Quasi-2D simulations of cell adhesion to wall surfaces under strong wetting affinity ($\gamma = -8$).** Time-resolved snapshots showing the effect of wetting phase volume ($\bar{\psi}$, top four rows; $\gamma = -8$ and $f_d = 0$) and shear flow ($f_d > 0$, bottom two rows; $\gamma = -8$ and $\psi = -0.5$) on particle capture by a neutral wall.
